# HaloClassifier: integrating coding-signature features and *k*-mer composition for plasmid-chromosome discrimination in haloarchaeal genomes

**DOI:** 10.64898/2026.09.21.753098

**Authors:** Lorena López-Roig, Alfred Fillol-Salom, Fernando González-Candelas

## Abstract

**Background:** Plasmids are key drivers of horizontal gene transfer (HGT), enabling the dissemination of accessory traits that shape microbial adaptation and ecological interactions. In *Haloarchaea*—dominant members of hypersaline environments—characterizing plasmidomes remains particularly challenging because most available genomes are incomplete, leaving many contigs unassigned to either chromosomal or plasmidic origin. This limitation hampers our ability to reconstruct archaeal plasmid diversity and to evaluate how plasmid-borne genes circulate within *Haloarchaea* and, potentially, outside this class.

**Results:** Here, we present HaloClassifier, the first machine-learning tool specifically designed to distinguish plasmid- and chromosome-derived contigs in haloarchaeal genomes. Unlike existing bacterial-focused approaches, HaloClassifier uniquely integrates haloarchaeal-specific genomic signatures with coding-derived features—an underused but highly informative class of predictors—and incorporates a variable-selection strategy that substantially reduces model complexity and computational cost while maintaining high predictive accuracy. When trained on simulated contigs generated from complete haloarchaeal genomes, the model achieves 96.61% accuracy on extralarge contigs (40–100 kb), increasing to 98.1% at a 0.6 classification threshold, with only 3.55% of contigs left unclassified. For small contigs (1–5 kb), the model achieves 79.51% accuracy overall and 84.42% at the 0.6 classification threshold, with 17.07% of contigs left unclassified. The model also maintains robust performance in metagenomic datasets.

**Conclusions:** By enabling reliable plasmid identification in fragmented assemblies, HaloClassifier fills a major gap and establishes the foundation for large-scale haloarchaeal plasmidome reconstruction. This framework will support future investigations into gene mobility, ecological adaptation, and the evolutionary dynamics of plasmids and HGT in *Haloarchaea*.

## Background

Plasmidomes—the complete repertoire of plasmids in an organism or microbial community—provide valuable insights into the mobility, distribution, and ecological significance of accessory genes [1]. By examining the plasmidome of a given environment, it becomes possible to track the flow of traits such as antimicrobial resistance, metabolic capabilities, or virulence factors, clarifying how these functions spread across lineages and shape microbial adaptation [2–5]. However, characterizing plasmidomes from high-throughput sequencing data remains challenging: both isolate-derived and, particularly, metagenomic assemblies frequently contain numerous contigs whose genomic origin (chromosomal or plasmidic) cannot be confidently resolved [6,7]. This ambiguity arises largely from assembly difficulties caused by repetitive regions, genome plasticity, uneven coverage, contamination, and the presence of minor populations masked by dominant taxa [7]. As a result, a considerable fraction of contigs remains unassigned, limiting our ability to interpret gene mobility and reconstruct the genetic structure of microbial communities.

To address this problem, a broad landscape of computational tools has emerged for classifying contigs as plasmid- or chromosome-derived. These methods—ranging from SVM-based models [8], Random Forests [9], and neural-network approaches [10] to graph-based approaches [11, 12] and, more recently, generative AI models [13]—have substantially improved our ability to recover plasmid content from fragmented assemblies. Yet, despite their diversity, all existing tools share a critical limitation: they were developed exclusively for bacteria, either targeting specific species (e.g., mlplasmids [8]) or providing domain-wide classifiers (e.g., PlasFlow [10], PlasClass [14]). Moreover, given that horizontal gene transfer (HGT) between bacteria and archaea has been demonstrated on multiple occasions [15–18], and that both domains coexist across a wide range of environments, focusing exclusively on bacterial plasmid classification limits our ability to obtain a broader view of HGT processes, particularly those involving archaea, and their contribution to microbial evolution.

*Haloarchaea*, the most dominant microorganisms in hypersaline environments [19], provide a particularly relevant system for studying gene mobility beyond bacterial systems, given their extensive HGT [17,20] and presence across diverse environments. Although primarily associated with extreme environments, *Haloarchaea* have also been detected in foods intended for human consumption [21,22] and in the human microbiota [23,24], suggesting that these microorganisms—and the mobile genetic elements they carry—may traverse ecological boundaries once considered prohibitive. However, despite their ecological relevance, the number of publicly available haloarchaeal genomes in databases such as GenBank [25] remains limited (fewer than 1,600 as of November 2025), and this number decreases by more than 80% (to 255) when considering only fully assembled genomes in which the genomic origin of each replicon is known. Together, these factors underscore the need for accurate haloarchaeal plasmid identification, both to illuminate gene flow within hypersaline ecosystems and to understand how plasmid-encoded traits may disseminate to distantly related taxa, potentially including clinically relevant bacteria.

To address this need, we introduce HaloClassifier, a Random Forest–based model specifically designed to classify contigs from partially assembled haloarchaeal genomes. Unlike existing bacterial plasmid classifiers, which primarily rely on sequence-derived genomic signatures (*k*-mer profiles and GC content), HaloClassifier also incorporates coding-derived features (codon usage, coding density, relative synonymous codon usage (RSCU), and effective number of codons (ENC))—which have been largely overlooked in plasmid classification—enabling a broader range of haloarchaea-specific signals to be captured. The pipeline incorporates a two-step feature-selection procedure that retains only the most informative predictors for the final model, substantially reducing computational cost and improving runtime efficiency. In addition, HaloClassifier is structured into four predefined contig size classes—small (1-5 kb), medium (5-10 kb), large (10-40 kb), and extralarge (40-100 kb)—allowing each size range to be modeled using the variable subset that best captures its distinctive chromosomal–plasmid signal.

Our results demonstrate that coding-derived features provide complementary information to sequence-derived features, significantly improving classification when combined with them. HaloClassifier achieves 96.61% accuracy on extralarge contigs (40–100 kb), increasing to 98.1% at a 0.6 classification threshold, with only 3.55% of contigs left unclassified. For small contigs (1–5 kb), the model achieves 79.51% accuracy overall and 84.42% at the 0.6 classification threshold, with 17.07% of contigs left unclassified. Importantly, the model enables, for the first time, large-scale plasmidome analysis in *Haloarchaea*, while its robust performance on metagenomic datasets extends its applicability to diverse environmental samples.

## Implementation

### Dataset Construction

To train and evaluate HaloClassifier, we compiled a high-quality dataset of haloarchaeal plasmids and chromosomes. Chromosomal sequences were obtained from all complete haloarchaeal genomes available in GenBank [25] and RefSeq [26] as of 24 November 2025, yielding 254 genomes (5 from GenBank and 249 from RefSeq). To ensure data quality and consistency, we retained only genomes with a single chromosome and at least one annotated plasmid, and filtered out assemblies showing ≥5% contamination or ≥25% strain heterogeneity as estimated with CheckM [27]. Additionally, chromosomes smaller than 1 Mb and genomes containing plasmids larger than 1 Mb were excluded to avoid potential misassemblies and inconsistencies in the annotation of megaplasmids.

Plasmid sequences were retrieved from PLSDB [28], a curated database that includes plasmids derived from both complete genomes and independent submissions, yielding 513 haloarchaeal plasmids. No additional quality filters were applied beyond excluding sequences larger than 1 Mb to minimize the inclusion of potential megaplasmids or misannotated secondary chromosomes.

After applying these criteria, 155 chromosomes and 510 plasmids were retained for downstream analyses. Most plasmids associated with the 155 selected complete genomes were also represented in PLSDB (353 of 422), while the remaining plasmids in the PLSDB collection expanded the dataset beyond the selected complete genomes. Given the relatively small size of the training dataset, we prioritized the use of the curated PLSDB collection to provide a consistent and quality-controlled source of plasmid sequences, although this may have resulted in the exclusion of some potentially suitable plasmids not represented in the final dataset. The complete lists of the training dataset and the corresponding genome quality assessment are provided in Supplementary Files S1–S3 in the Zenodo repository [10.5281/zenodo.22233815].

### Fragment generation across four size ranges

To improve classification performance across heterogeneous contig sizes, we adopted a multi-length training strategy inspired by PlasClass [14], with a key modification for haloarchaeal replicons. Instead of training on fixed fragment lengths, we defined four size ranges: Small (1–5 kb), Medium (5–10 kb), Large (10–40 kb), and Extralarge (40–100 kb), and trained separate models on variable-length fragments within each range.

Within each size class, plasmid and chromosomal sequences were randomly fragmented to generate representative and non-redundant datasets. Fragmentation was performed on plasmids first, as they often limit the number of non-redundant fragments that could be generated—particularly in the larger size ranges—, and chromosomal fragments were subsequently sampled to approximately match plasmid counts. Fragmentation parameters (sampling depth and allowed overlap) were adjusted per size range to compensate for the scarcity of long plasmids while minimizing redundancy.

To avoid oversampling small plasmids, depth amplification was applied only to plasmids exceeding a minimum size threshold, defined as four times the average fragment length of each range (e.g., ≥100 kb for Large and ≥280 kb for Extralarge). An exception was made for the Medium range, where a threshold of ≥40 kb was sufficient due to higher plasmid availability.

Plasmid replicons were fragmented sequentially using random fragment sizes within each interval, with a baseline overlap of 30% (increased to 70% in the Extralarge range). Depth amplification was applied to all ranges except Small, and only to plasmids above the minimum size threshold in each case. For the Medium range, plasmids were sampled at an additional depth of 2, while Large and Extralarge plasmids used a depth of 4. To limit redundancy among depth-amplified fragments, the maximum allowed overlap for these additional passes was reduced to 15%, except in the Extralarge range, where a 60% threshold was required due to the scarcity of long plasmids.

Chromosomal replicons were sampled differently: instead of sequential fragmentation, fragments were drawn from random genomic positions to approximately match plasmid fragment counts. Overlap between chromosomal fragments was strictly controlled and depended on size range: no overlap was allowed in the Small range, while maximum overlaps of 60% (Medium) and 70% (Large and Extralarge) were permitted in the larger size bins.

This procedure yielded balanced datasets of plasmid and chromosomal fragments across all size ranges, with fragment counts summarized in **Table 1**.

**Table 1.** Number of fragments obtained from the training dataset across four size ranges. : Small (1–5 kb), Medium (5–10 kb), Large (10–40 kb), and Extralarge (40–100 kb). For the Extralarge model, up to 96 fragments per chromosome were sampled; however, shorter genomes occasionally yielded slightly fewer fragments (94–95), resulting in minor deviations from the expected total.

| Size range | Plasmid fragments | Chromosome fragments |
| --- | --- | --- |
| Small (1-5 kb) | 55,912 | 55,800 (360 p/chr) |
| Medium (5-10 kb) | 54,373 | 54,250 (350 p/chr) |
| Large (10-40 kb) | 26,650 | 26,660 (172 p/chr) |
| Extralarge (40-100 kb) | 14,563 | 14,623 (96 p/chr) |

### Optimum-feature selection

For each fragment, we computed two complementary families of features capturing both genomic compositional signals and coding-related properties (Table 2). Fragment length was included as a control (confounding) variable. All *k*-mer profiles were expressed as relative frequencies normalized by the total number of *k*-mers in the fragment.

**Table 2.**
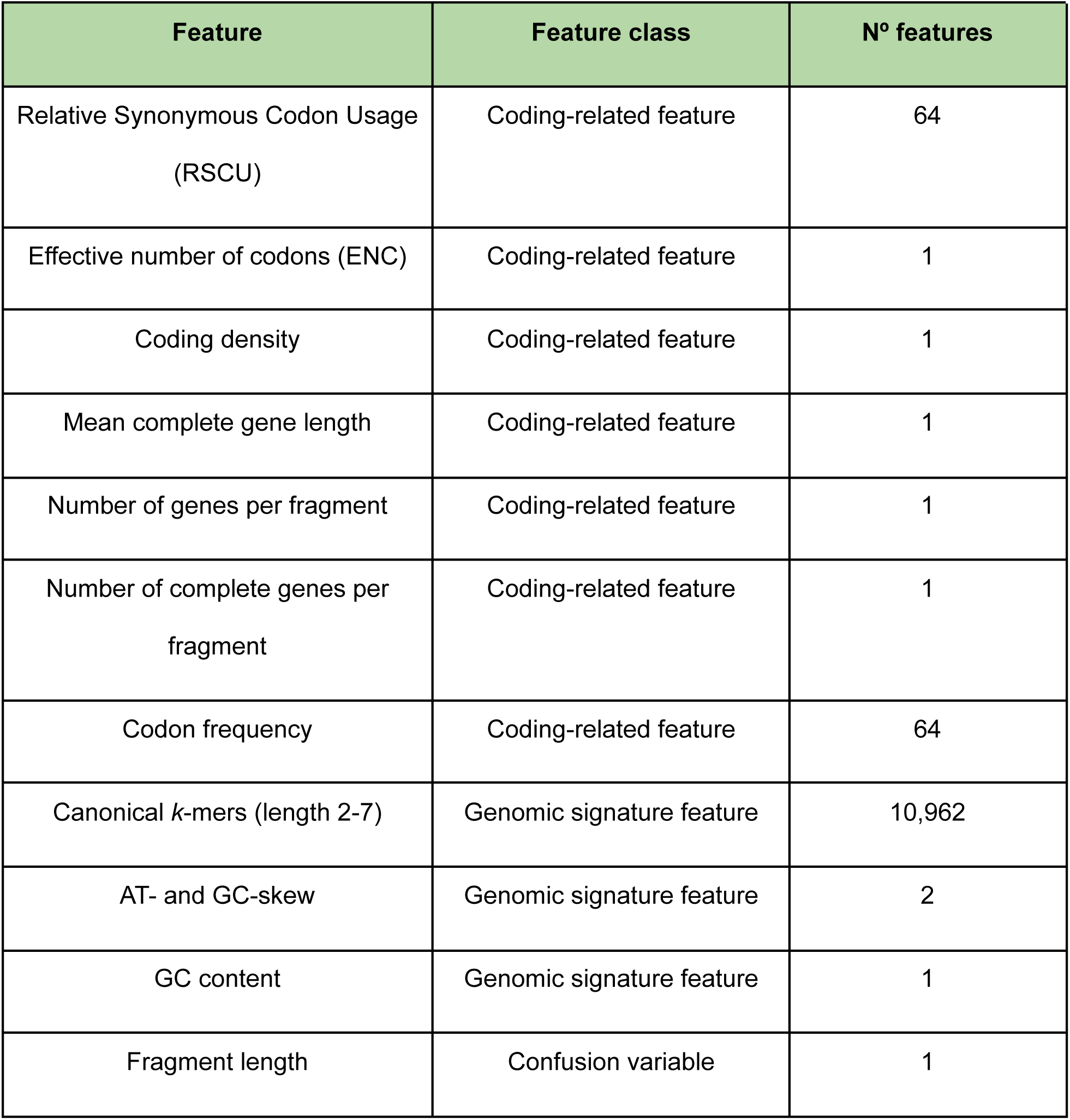
Summary of all features computed for each fragment, grouped into coding-related, genomic signature, and confusion-variable classes. *K*-mer profiles and codon frequencies correspond to relative frequencies normalized by fragment length and total codon count, respectively.

Coding-related features were derived from gene predictions obtained with Prodigal [29] in metagenomic mode. Codon-usage statistics were computed from both complete and partial coding sequences (CDSs), because excluding truncated genes would substantially reduce the available codon signal, particularly in short fragments. For partial CDSs, trailing nucleotides at the 3′ end were trimmed when necessary to preserve the reading frame, and codon frequencies were normalized to sum to 1 per fragment. Each codon frequency was retained as a separate feature in the feature matrix, resulting in 64 codon-frequency features per fragment.

A total of 11,099 features were obtained, including 133 coding and 10,965 non-coding features, in addition to the contig length control variable. To improve prediction performance and adapt the models to size range, an optimal feature subset was selected independently for each predefined class (Small, Medium, Large and Extralarge). This size-specific feature selection allows different variables to contribute differently depending on fragment length, enhancing both model specificity and overall predictive accuracy. Importantly, such reduction in dimensionality is not only computationally advantageous: given the relatively limited number of complete haloarchaeal genomes available for training, retaining all variables would disproportionately increase the risk of overfitting. By restricting the model to the most informative features, we ensure stable performance even under data-sparse conditions, while simultaneously reducing model complexity and computational cost—an essential requirement for the large-scale analyses typical of modern metagenomics.

This approach ensures that, although some non-informative variables (e.g., RSCU values for stop codons or for codons encoding methionine and tryptophan, which are represented by a single codon) and variables with potentially noisy signals, particularly in shorter fragments, are initially included, the final model is trained using a reduced subset of predictors selected to achieve high predictive accuracy while minimizing model complexity. To achieve this, a two-step procedure was implemented:

Step I – Coarse feature selection: The procedure began with an initial set of 10 variables (as models with zero predictors cannot be trained), and variables were subsequently added in batches of 30 according to their Random Forest importance scores. For each variable count, 25 repetitions were performed, and mean accuracy and standard deviation were recorded. The 1-standard-error rule was then applied, selecting the smallest set of variables whose accuracy fell within one standard deviation of the maximum.

Step II – Fine feature selection: Starting from the variable count obtained in Step I, an interval of ±50 variables was explored in increments of 5. Each configuration was repeated five times, and the final number of variables was selected by maximizing mean accuracy after subtracting the standard deviation. In the case of ties, the most parsimonious model was selected. It should be noted, however, that there is no unique “optimal” number of variables, as this may vary depending on the study objective or the nature of the microorganisms analyzed.

All repetitions were performed using fixed random seeds to ensure reproducibility.

To prevent unnecessary computational costs during the coarse feature-selection phase, an early-stopping criterion was implemented (Figure S1). Specifically, the algorithm was configured to terminate if, across ten consecutive steps (each adding 30 new variables), none of the evaluated models exceeded the highest accuracy observed up to that point. This criterion was designed to identify *plateau* regions in the feature-accuracy relationship, where further increases in the number of variables were unlikely to provide meaningful improvements in predictive performance. Continuing beyond this point would increase model complexity and training time without an expected practical gain in discriminative performance. The script for this feature-selection strategy is available as File S4.

### Building the machine-learning model

Random Forest classifiers were trained independently for each fragment size range using the subsets of features selected in the previous steps. Random Forest was chosen because it handles high-dimensional, heterogeneous genomic features robustly, is resistant to overfitting, and performs well with limited training data [30]—conditions that match the characteristics of our dataset. All models were trained with 100 trees (n_estimators = 100), which provided a good balance between computational cost and predictive accuracy. For each size range, the fragments obtained were split randomly into training and test sets (80/20), and model performance was evaluated using accuracy, precision, recall, F1-score and Matthews correlation coefficient (MCC).

### Statistical analysis

To evaluate the robustness of HaloClassifier, 25 independent genome-fragmentation replicates (N1, N2…; N25) were generated for each size range using the same criteria described above (overlap and read depth). Performance metrics (accuracy, precision, recall, F1-score, and MCC) were computed for each replicate and for each class (chromosome and plasmid) when applicable. Bootstrap tests [31] were then performed to assess whether the final model’s performance differed significantly from the distribution of replicate results.

To assess the contribution of coding features to classification accuracy, three complementary model configurations were evaluated: (i) the full model including all selected variables (Final Model); (ii) a model trained exclusively on coding features (Only Coding); and (iii) a model excluding all coding-related variables (No Coding). In both reduced-feature configurations, fragment length was included as a confounding variable. Paired Wilcoxon signed-rank tests [32] were used to compare the performance of each reduced-feature model against the Final Model across all contig-length categories.

### Comparison with other softwares

To evaluate the performance of HaloClassifier, we compared it with PlasForest [9], PlasFlow [10], and PlasClass [14], tools that were originally trained on bacterial genomes. Since no dedicated plasmid–chromosome classifier exists for archaeal data, these tools were included as a baseline reference for cross-domain performance.

For this benchmark, we randomly selected 1,000 fragments per size category (500 chromosomal and 500 plasmid-derived) from a single bootstrap replicate (N14) of our evaluation dataset. These fragments were not used during the training of HaloClassifier, although such inclusion should not have influenced performance. In total, 4,000 simulated contigs, spanning four length ranges, were used to construct realistic prediction scenarios and enable a fair comparison across tools.

### Metagenomic dataset and evaluation strategy

To evaluate the applicability of HaloClassifier to real metagenomic data, we analysed its predictions across 233 metagenome-assembled genomes (MAGs) belonging to the class *Halobacteria* from the GEM dataset [33], downloaded on 3 March 2026. The dataset comprises a wide range of curated environmentally derived genomes. Among these, 57 MAGs originated from laboratory enrichment cultures grown in defined media, whereas the remaining 176 correspond to genomes assembled directly from environmental metagenomic samples. Given potential differences in assembly quality and genomic composition between these sources, analyses were performed separately for both groups. The complete lists of environmental and enrichment-derived MAGs included in this study, together with their associated metadata, are provided in Supplementary Files S5 and S6.

To assess model performance in a genome-wide context, we quantified, for each MAG, the proportion of its total assembled sequence classified above each confidence threshold (≥0.6, ≥0.7, and ≥0.8). To assess the relationship between genomic relatedness to the training dataset and classification performance, we calculated pairwise Average Nucleotide Identity (ANI) using FastANI [34], comparing each MAG against all genomes in the training dataset. For each MAG, the genome yielding the highest ANI was retained as the best-matching reference, and its ANI value was compared with the proportion of MAG sequence classified at the highest confidence threshold (≥0.8). This analysis was performed separately for environmental and enriched MAGs.

As an additional analysis, we assessed whether the relationship between genomic relatedness and classification performance was also reflected in broader taxonomic relationships. For each MAG, the taxonomic assignment of the MAG was compared with that of its best-matching training genome. MAGs were assigned to three categories: same genus, same family (different genus), and different family. Taxonomic relationships were determined using a hierarchical, conservative scheme that prioritised genus-level information. MAGs were classified as same genus when both genera were defined and identical, and as same family or different family when both genera were defined but differed and the corresponding families were the same or different, respectively. When one or both genera were undefined, a same genus or same family relationship could not be asserted if the families were identical, because the undefined genus could correspond either to the same or a different genus; such cases were therefore conservatively assigned to unknown. When the families differed, these cases were classified as different family. This approach avoided inferring taxonomic relatedness from incomplete assignments. Taxonomic assignments were reviewed and, where necessary, updated according to the current NCBI Taxonomy classification prior to this analysis. All taxonomic corrections, including revised family assignments and genus-level labels that were reclassified as undefined, are provided in Supplementary File S7.

### Pipeline description

HaloClassifier is a Python package installable via pip, designed to classify haloarchaeal contigs as chromosomal or plasmidic using size-specific machine-learning models trained on haloarchaeal genomes. The workflow consists of two integrated command-line tools that perform feature extraction and classification, corresponding to the steps summarized in Figure 1.

**Figure 1.**
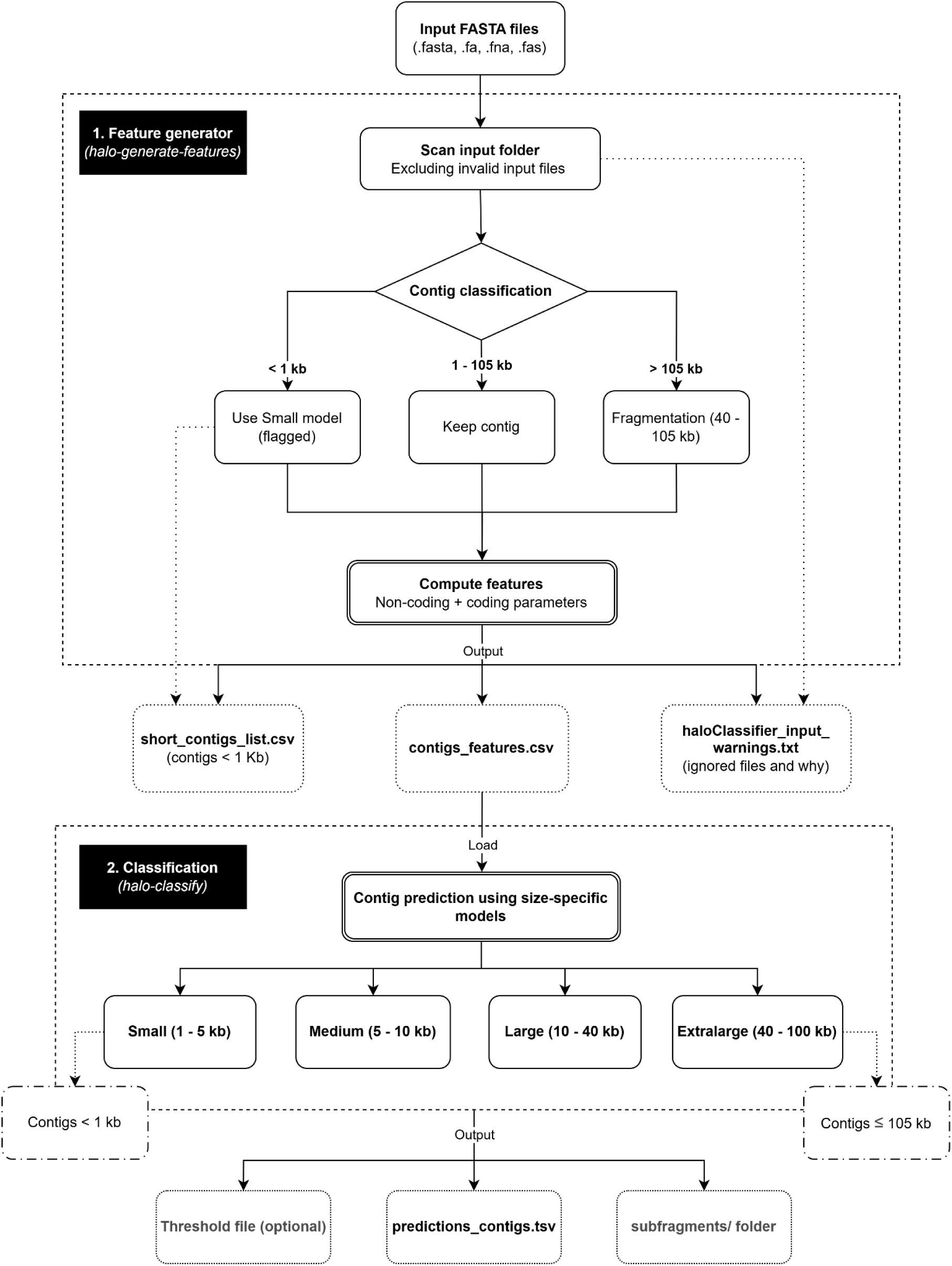
**Two-step workflow of HaloClassifier**, consisting of feature generation and subsequent contig classification using size-specific models. Diagram created with draw.io.

The pipeline begins by assigning each input contig to one of the size classes used during model training based on its fragment length. This ensures that every sequence is evaluated with the model optimized for its size range, preventing model mismatch and improving prediction accuracy. Contigs <1 kb are processed under the Small model but flagged as being outside the training range. Contigs >105 kb are automatically fragmented into pieces between 40 kb and 105 kb. The additional 5 kb tolerance avoids unnecessary fragmentation of near-threshold contigs while maintaining compatibility with the Extralarge model.

Feature extraction is performed with the command halo-generate-features, which first assigns each contig to its size class (small, medium, large or extralarge) and then computes only the subset of sequence-derived features required by the machine-learning model corresponding to that class. The output is a feature matrix in CSV format, and it can be accompanied by a file of flagged contigs (e.g., <1 kb) and warnings for ignored files if applicable. This step supports multithreading (e.g., -t 8) and can process multi-contig FASTA files.

Classification is performed with the command halo-classify, which loads the appropriate pre-trained model for each contig or fragment. Predictions include the assigned label (chromosome or plasmid) and the corresponding probability scores. For contigs that require fragmentation, the final output reports the mean probability across all fragments, while fragment-level predictions are saved in a separate folder (*subfragments/*). Users can optionally specify a probability threshold to retain only high-confidence predictions (e.g., -p 0.6), in which case an additional filtered report is created.

This program is available in GitHub: https://github.com/Lorena0105/HaloClassifier

To evaluate computational performance, we randomly selected 1,000 Extralarge contigs (40–100 kb) from the N14 robustness replicate (see above). Runtime and peak memory usage were measured over 10 independent executions using a single CPU thread. HaloClassifier completed the analysis in an average of 316.9 s, with a peak memory usage of approximately 210 MB. Most of the execution time was spent on the feature extraction stage (309.2 s), whereas the classification step required only 7.7 s on average, with a peak memory usage of approximately 176 MB. Under the same conditions, PlasForest completed the analysis in an average of 320.2 s with a peak memory usage of 1.18 GB. The effect of sequence size and thread count on runtime and peak memory usage for both tools is shown in Figure S2.

All computational benchmarks were performed on compute nodes providing 64 physical cores (128 logical CPUs) and 732 GB of RAM.

## Results

### Taxonomic and architectural diversity of the training dataset

Given that HaloClassifier was trained on a relatively small number of complete genomes (155 chromosomes) and an independent set of plasmids (510 sequences from PLSDB), we first assessed whether the training dataset adequately captured the taxonomic and replicon diversity currently represented within *Haloarchaea*. To this end, we evaluated the taxonomic distribution of both the chromosomal and plasmid components separately (Figure 2).

**Figure 2.**
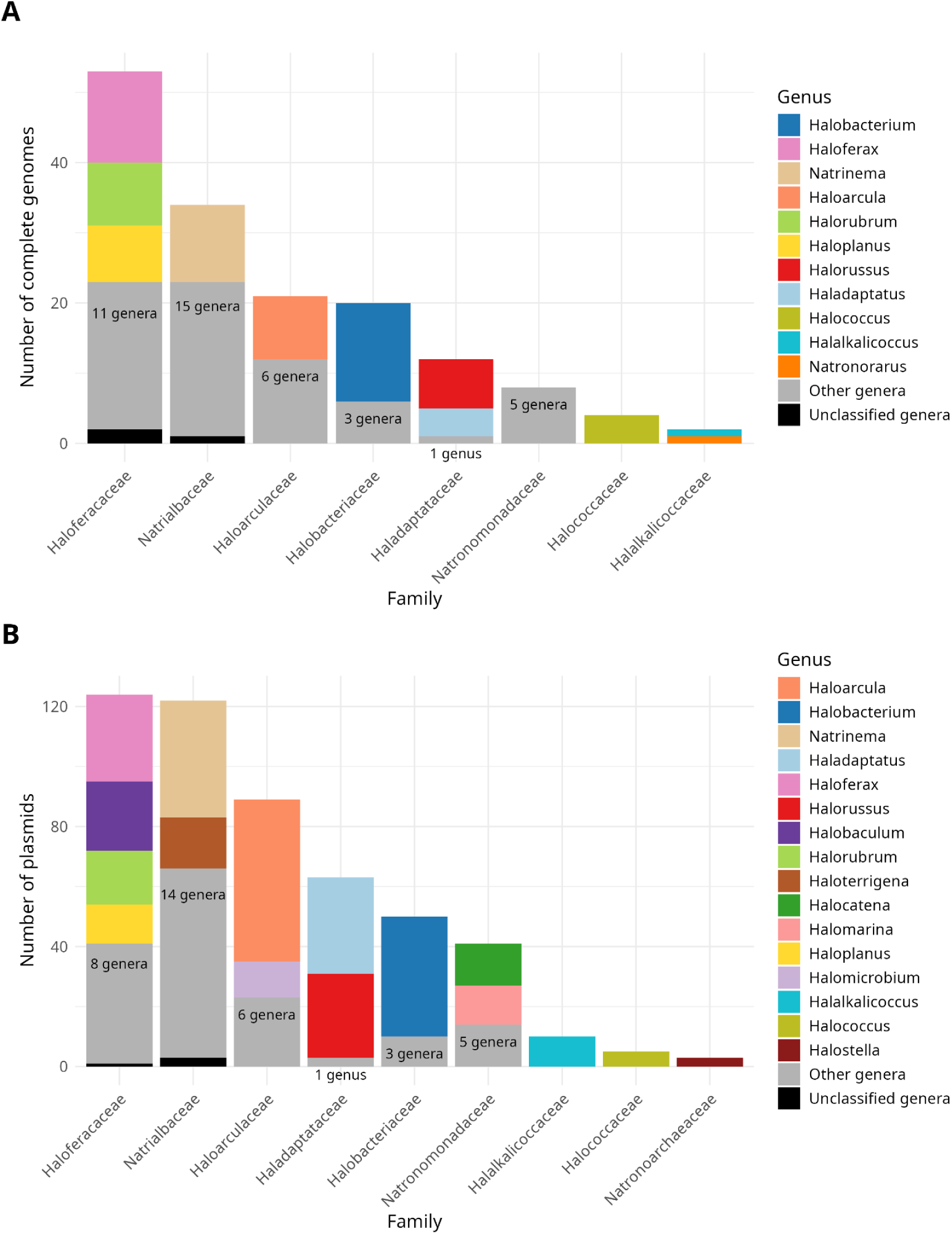
Taxonomic distribution of the training dataset. (A) Family-level distribution of the 155 chromosomal sequences derived from complete haloarchaeal genomes. Genera were highlighted if they were the sole genus within a family, represented >30% of the family-level counts, or comprised >5 complete genomes. (B) Family-level distribution of the 510 plasmid sequences retrieved from PLSDB. Genera were highlighted using the same criteria, except that the abundance threshold was set to >10 plasmids per genus. In both panels, highlighted genera are shown with distinct colors, consistent across panels where applicable. The categories “Other genera” (grey) and “Unclassified genera” (black) group remaining taxa; the number within the “Other genera” segment indicates how many genera it includes.

Of the eleven families currently described within the class *Haloarchaea*, eight contain at least one complete genome that meets the quality criteria defined for this study (see Methods for more details). However, four families—*Haloferaceae*, *Natrialbaceae*, *Haloarculaceae*, and *Halobacteriaceae*—account for 85% of all currently available complete genomes (Figure 2A), resulting in an overrepresentation of these taxa in the training set. Within these families, a small number of genera also dominates the dataset, whereas the remaining genera contribute only marginally; these dominant genera are highlighted with distinct colors in Figure 2.

In contrast, the taxonomic distribution of the plasmid dataset (Figure 2B), which includes sequences from both complete genomes and independent PLSDB submissions, shows a broader and slightly more balanced representation across families, although the same general pattern of dominance by a limited number of families is maintained. Notably, the plasmid dataset includes five genera that are not represented in the chromosomal dataset—*Halostella*, *Halobiforma*, *Halosegnis*, *Halovenus* and *Salinirubellus*—as well as an additional family, *Natronoarchaeaceae*, represented by *Halostella*. These taxa expand the taxonomic diversity captured by the training dataset beyond that provided by complete chromosomes alone. Conversely, four genera represented among chromosomes—*Halorarum*, *Halovalidus*, *Natronorarus*, and *Salinirarus*—are absent from the plasmid dataset.

Beyond taxonomic diversity, we also examined the plasmid content and architecture of the complete genomes represented in the training dataset. Although plasmid sequences were retrieved from the curated PLSDB collection [28], most plasmids present in the 155 selected completed genomes were also represented in PLSDB (353 of 422), providing a strong correspondence between the plasmid component of the training dataset and the plasmid complement observed in complete haloarchaeal genomes (see Implementation).

On average, the *Haloarchaea* genomes included in the training dataset contained 2.72 plasmids per assembly (standard deviation: 1.53; range: 1–7). Plasmid size was even more variable, with a mean of 221.1 kb (standard deviation: 170.6 kb; range: 1.4–916 kb), indicating that haloarchaeal plasmids span a broad size spectrum. Consequently, plasmid content also varied substantially among genomes. As shown in Figure S3, plasmid content ranged from 2.53% to 31.81% of the total genomic DNA across the best-represented genera, with considerable variation observed not only among genera but also among strains of the same species. For example, *Haloferax* displayed relatively consistent plasmid content (17.54–29.03%), whereas *Haloarcula* and *Natrinema* exhibited much broader ranges (3.12–31.81% and 2.53–29.17%, respectively), emphasizing the diversity of plasmid architectures represented in *Haloarchaea*.

Together, these results indicate that, despite the limited number of complete haloarchaeal genomes currently available, the training dataset captures substantial taxonomic and plasmid diversity. In particular, the high variability in plasmid number, size, and relative genomic contribution highlights the importance of plasmids as a substantial and heterogeneous component of haloarchaeal genomes, further supporting the need for accurate chromosome–plasmid classification.

### Reliability of the classification method

Given that 80% of the fragments in the training dataset were allocated to model training, reshuffling the original fragments would produce training sets that differ only marginally from one another. To more rigorously assess the robustness of HaloClassifier, we therefore generated 25 independent genome-fragmentation replicates for each size range from the same complete genomes, using the original overlap and read-depth criteria (see Implementation). Performance metrics were computed for each replicate and compared with those obtained for the final model. Across the four contig-size categories, the performance of the final model closely matched the distribution of the 25 replicates (Figure 3).

**Figure 3.**
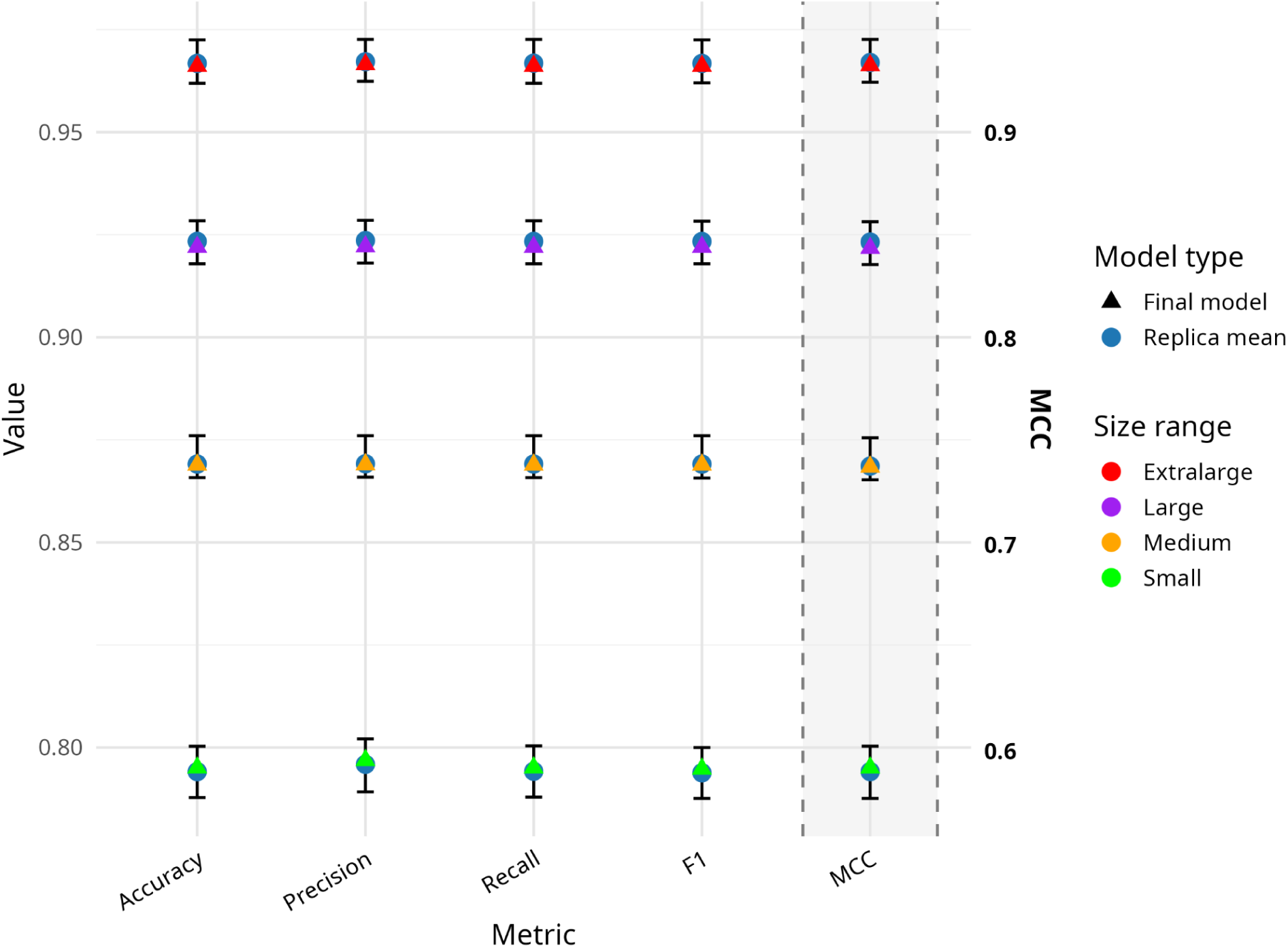
Robustness of model performance across fragment size ranges. Colored triangles indicate the final model’s value for each metric (Accuracy, Precision, Recall, F1, MCC) across fragment sizes (green = Small, orange = Medium, purple = Large, red = Extralarge). Dark blue circles show the mean value of 25 independently generated fragment sets, with error bars representing the observed minimum and maximum. MCC values, shown on a separate scale on the right to avoid compressing the range of the other metrics, are highlighted with a shaded region and displayed in bold. Bootstrap tests were performed to evaluate whether the performance of the final model differed significantly from that obtained using independently generated fragment sets (Table 3). Across all fragment sizes and evaluation metrics, the bootstrap *p*-values were consistently high (all > 0.60), indicating no statistically significant differences between the final model and the distribution of replicate results. This demonstrates that the model’s performance does not depend on any particular fragmentation instance of the genomes.

**Table 3.**
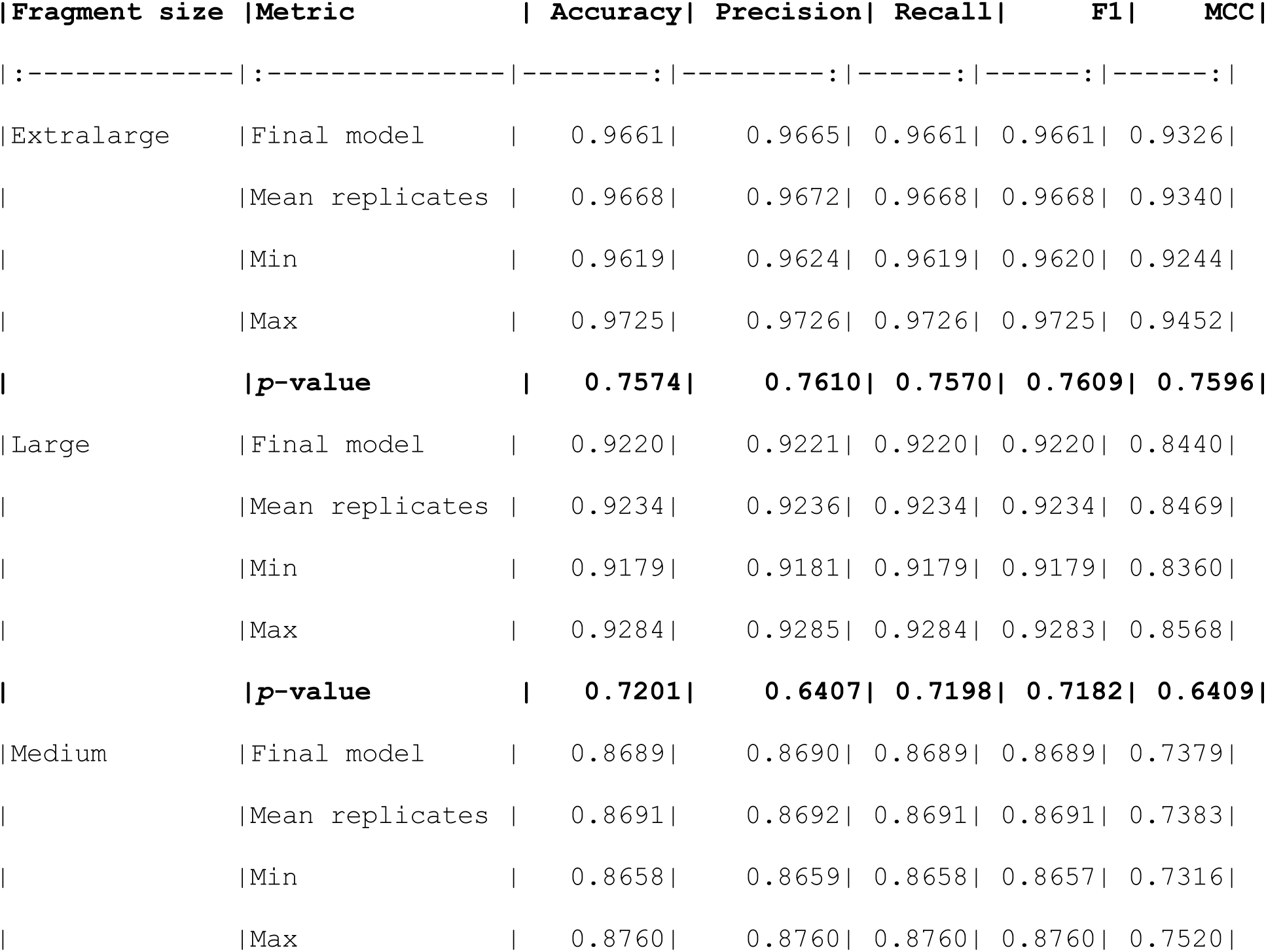
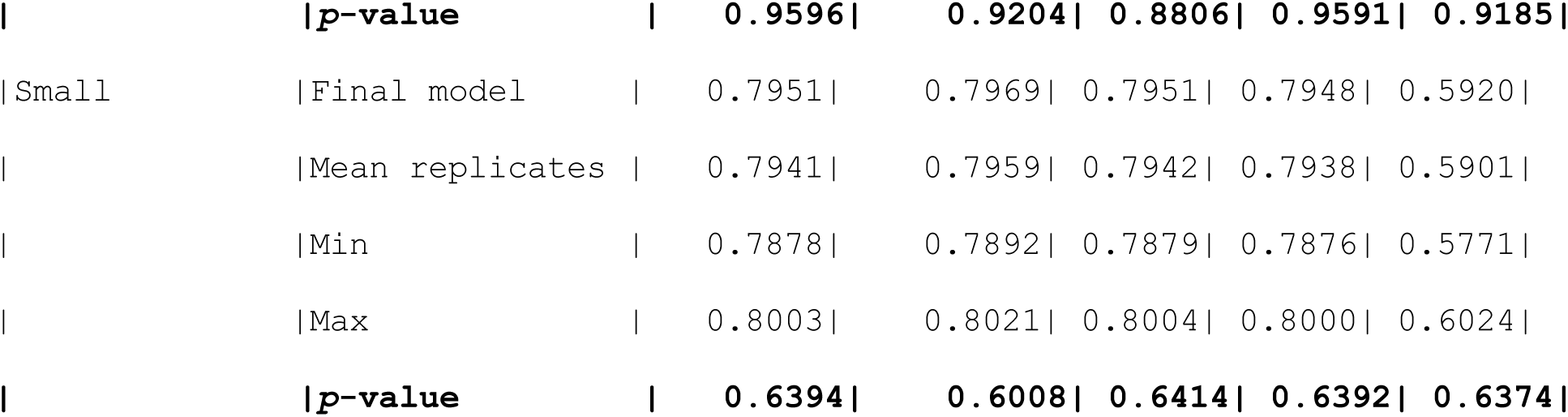
Bootstrap summary for each fragment size. Comparison between the final model and 25 independently generated replicate sets for each metric (Accuracy, Precision, Recall, F1, MCC). The table reports the final model performance, the mean, minimum, and maximum values observed across replicates, and the *p*-value from the bootstrap significance test.

The performance of the final model closely matched the mean performance of the 25 replicate sets, typically differing by less than 0.002 in absolute terms in all parameters. The narrow ranges defined by the minimum and maximum replicate values further indicate that alternative fragmentations introduce only minimal variability into the evaluation. Importantly, even for the smallest fragments—where performance variability is usually greatest—the final model remained tightly aligned with the replicate distributions.

Together, these results show that HaloClassifier behaves consistently across independent fragmentations of the same genomes. This robustness indicates that the classifier is resilient to realistic variation in contig boundaries and that its reported performance metrics are representative of the underlying data rather than a consequence of a specific training fragmentation.

### Coding features play an increasingly important role in the classification of smaller contigs

Short contigs pose a particular challenge for genomic classification, as their limited sequence context can introduce biases that disproportionately affect *k*-mer–based measures—the type of features most commonly used for this task—which can ultimately compromise classification accuracy. As shown in Figure 4A, the relative importance of coding-derived features increases as contig length decreases, reaching 25% and 26% in the Medium and Small models, respectively. Conversely, in the Extralarge model, coding-derived features account for 11% of the total feature importance, indicating a greater contribution from sequence-composition features when more sequence context is available. When focusing on the 100 most important variables (Figure 4B), the relative contribution of coding-derived features increases across all size ranges, especially in the Extralarge model. Thus, coding-derived features are not only retained in the final feature sets but are also slightly enriched among the highest-impact predictors.

**Figure 4.**
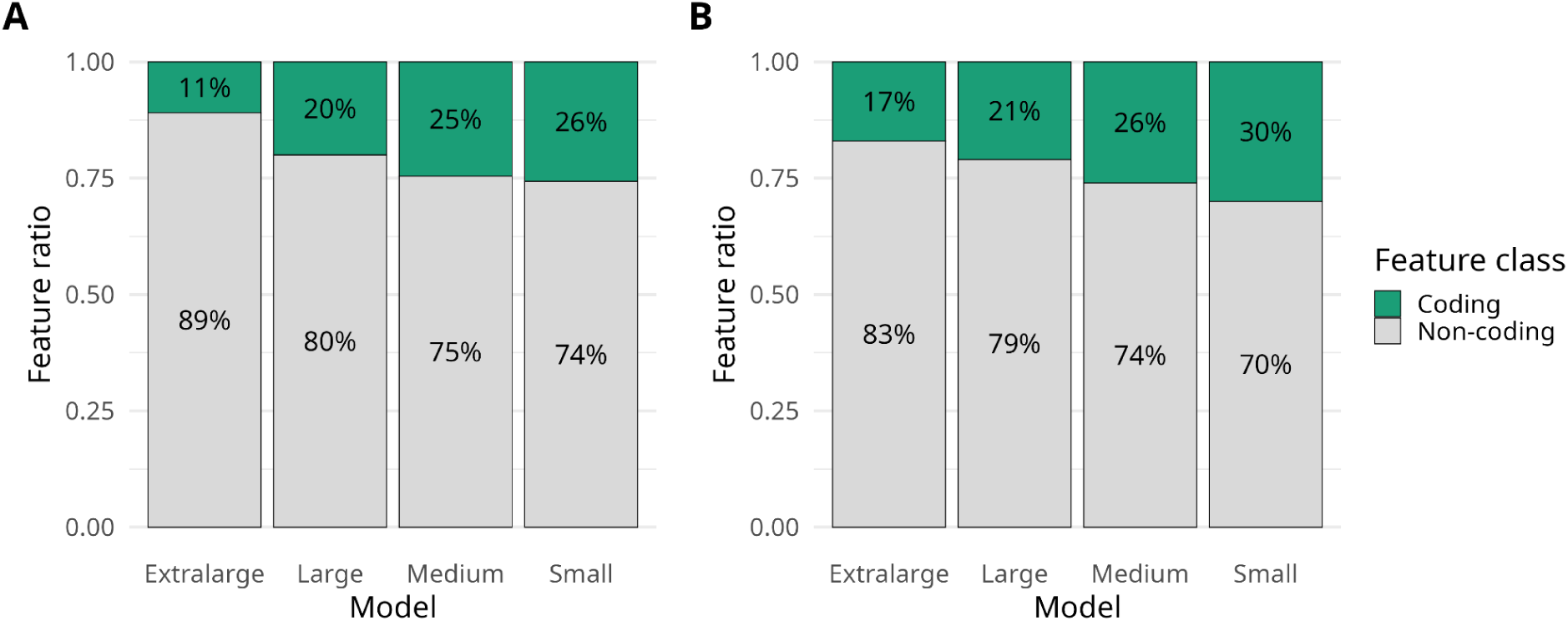
Relative contribution of coding (green) and non-coding (grey) variables across the size categories defined in HaloClassifier. Panel A shows the complete set of predictors retained after the fine phase of feature selection (Step II), representing the optimal subset that maximizes mean accuracy. Panel B displays the top 100 variables within this same optimal subset, ranked by their Random Forest feature-importance scores during model training.

The feature-selection procedure yielded subsets of 265, 350, 355, and 595 variables for the Small, Medium, Large, and Extralarge models, respectively. This increase in the number of retained variables with contig length suggests that longer fragments provide a broader range of predictive signals that can be exploited by the model. The complete outputs of the feature-selection procedure are provided in the Supplementary Material. Summary statistics and individual outcomes from the coarse feature-selection step across the four size-specific models are reported in Files S8–S15, while the complete ranked lists of features retained after fine feature selection, together with their corresponding Random Forest importance scores, are provided in Files S16–S19.

Among the coding features, the codon GCC, which encodes alanine, consistently appears as a highly important variable across all size categories. In the Extralarge model, *codon_GCC* ranks first and *RSCU_GCC* second, while in the Large model they rank first and third, respectively. For Medium contigs, *codon_GCC* remains in the top 5 (rank 4), although *RSCU_GCC* drops to rank 24, and for Small contigs, *codon_GCC* still resides within the top 10 (rank 7), whereas *RSCU_GCC* drops further (rank 35), reflecting the increased noise in smaller fragments. Remarkably, *codon_GCC* maintains a top-10 position across all size ranges, indicating that alanine codon usage constitutes a stable and recurrent discriminatory feature between chromosomal and plasmid-derived contigs in *Haloarchaea*. Figure S4 presents the top 10 variables for each size category.

To quantify the relative impact of coding variables on classification performance, we compared three complementary model configurations: (i) the Final Model, which uses the complete set of selected variables; (ii) Only Coding, trained using exclusively coding-derived variables; and (iii) No Coding, in which all coding-related predictors were removed. In both reduced-feature configurations, fragment length was included as a confounding variable. To make the comparison robust and directly comparable across configurations, we reused the 25 independent fragmentations generated in the robustness analysis. For each replicate and contig-length category, we applied the same two-step feature-selection protocol used for the Final Model, allowing each run to identify its own optimal subset of predictors.

Performance across the three configurations was evaluated using paired Wilcoxon signed-rank tests on the 25 matched replicates. Figure 5 summarizes the differences between each reduced-feature configuration and the Final Model for the Small (top) and Extralarge (bottom) contig-length models. The complete performance values (mean ± SD), together with the corresponding paired Wilcoxon signed-rank test *p*-values, are provided in Table S1.

**Figure 5.**
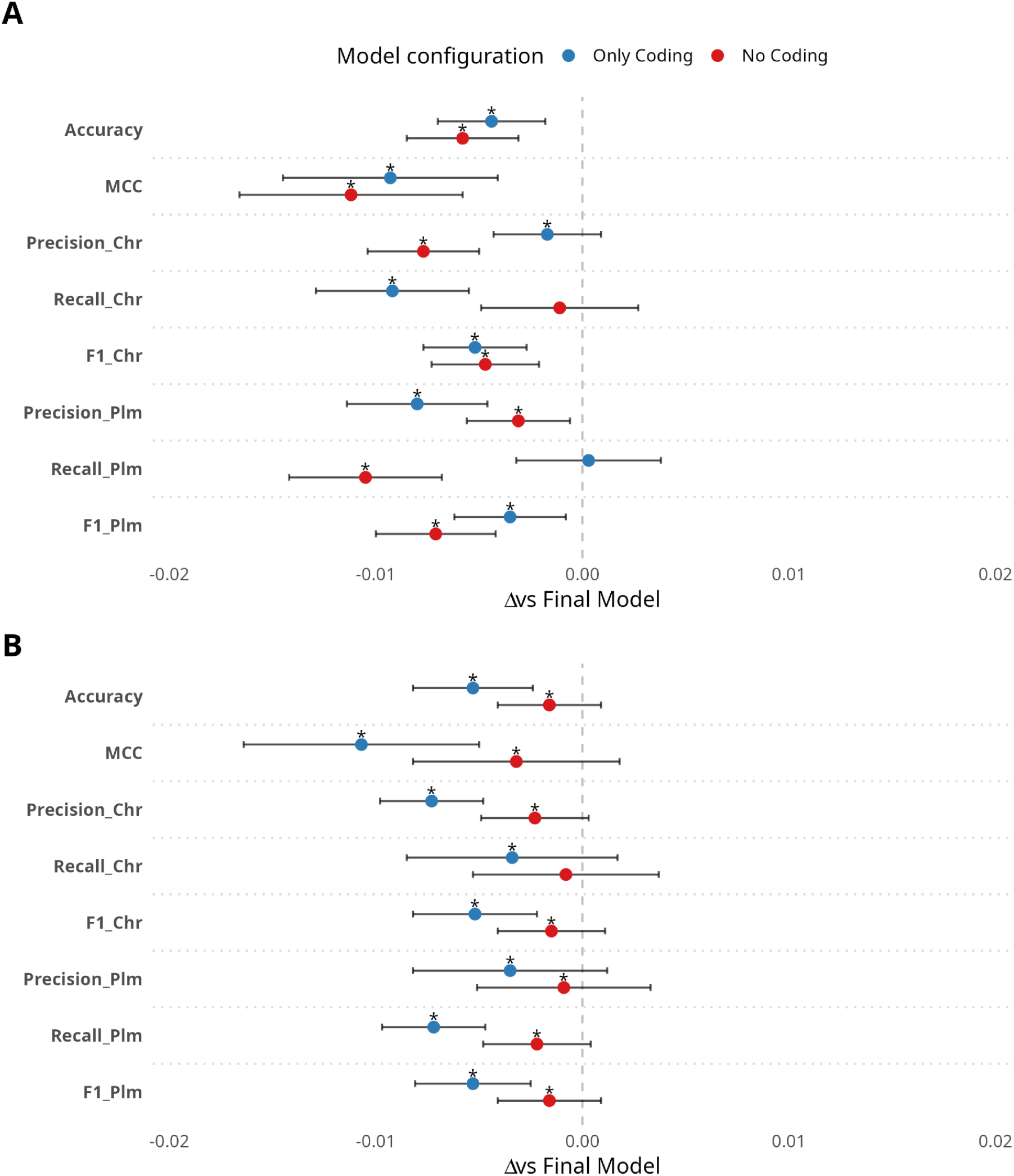
Effect of feature removal on model performance. Comparison of two reduced configurations—*Only Coding* (blue) and *No Coding* (red)—against the full-feature Final Model in Small (A) and Extralarge (B) models. The Final Model includes all available features. Points show mean performance per metric, and error bars indicate standard deviation. Asterisks denote statistically significant differences from the Final Model based on paired Wilcoxon signed-rank tests (*p* < 0.05). Exact *p*-values are reported in Table S1.

Both reduced-feature configurations exhibit statistically significant decreases relative to the full model in most metrics; however, these drops remain small in practical terms—typically below 1%—indicating that much of the discriminatory signal captured by the Final Model is preserved even when coding or non-coding feature sets are considered independently. This effect is particularly notable for the Only Coding model, which is constrained to a maximum of 133 features, whereas the No Coding model is selected from a substantially larger candidate feature space (∼11,000 variables) through an optimal feature selection procedure.

Notably, as contig size increases, the performance gap between the two reduced-feature configurations becomes more pronounced: although Only Coding and No Coding perform similarly for Small and Medium models, No Coding progressively outperforms Only Coding in the Large and Extralarge models. This trend shows that smaller contigs rely more on coding-derived information, whereas in longer sequences the non-coding *k*-mer signal becomes more stable, more representative of the genomic context, and, ultimately, more informative. However, it should be noted that when both sets of variables (coding and non-coding) are considered together, performance is consistently higher across almost all metrics, indicating that the two feature blocks provide complementary sources of information.

### Performance of alternative plasmid identification tools on haloarchaeal contigs

To benchmark HaloClassifier against other methods designed for the same purpose, we evaluated three widely used plasmid identification programs. Since no plasmid identification tools have been specifically trained on archaeal or haloarchaeal genomes, we compared our method with tools trained exclusively on bacterial data—PlasForest [9], PlasFlow [10], and PlasClass [14]—as a reference point.

PlasForest, which relies on homology searches against a pre-compiled bacterial genome database, proved unsuitable for haloarchaeal genomes. As haloarchaeal sequences are entirely absent from its reference dataset, the method classified all 4,000 fragments as chromosomal. Furthermore, PlasForest had a substantially longer runtime than PlasFlow and PlasClass.

By contrast, PlasFlow and PlasClass—both *k*-mer–based methods designed to capture the wide diversity of bacterial sequences—produced meaningful predictions that could be compared with HaloClassifier. For these tools, we applied a probability threshold of 0.6, leaving fragments within the 0.4–0.6 range unclassified. Figure 6 summarizes the main performance metrics under this threshold for the Extralarge model (the rest can be found in Figure S5). The percentage of unclassified fragments is also shown. When available, per-class metrics (chromosome vs plasmid) are reported to highlight whether each method exhibits systematic biases toward either genomic origin. The corresponding confusion matrices for all tools and contig-length categories are provided in Table S2. The effect of different confidence thresholds on HaloClassifier performance across all contig-length categories is summarized in Table S3.

**Figure 6.**
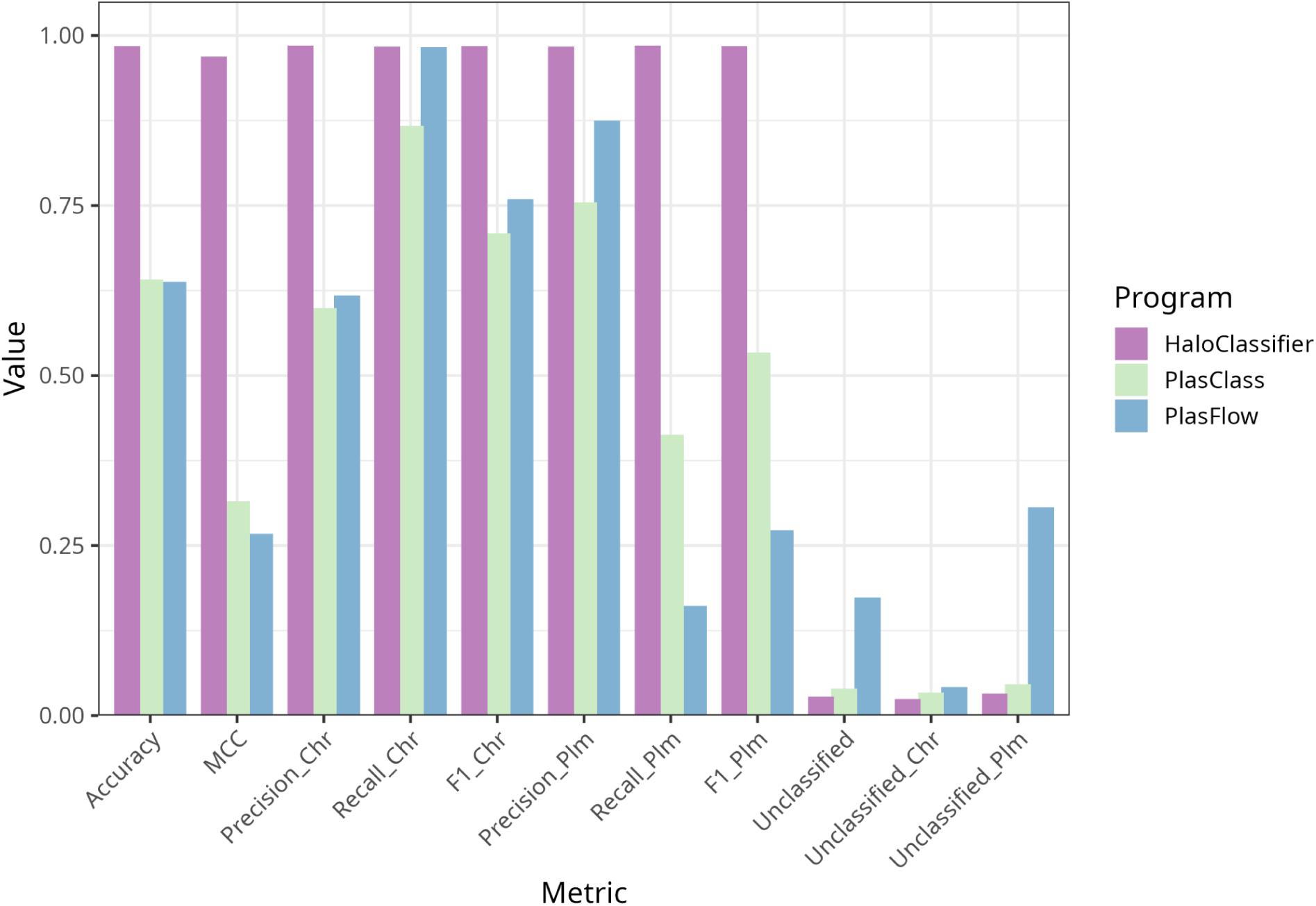
Performance comparison of three plasmid identification tools on simulated contigs from complete haloarchaeal genomes (Extralarge model). PlasClass (green) and PlasFlow (blue) were trained on bacterial genomes, whereas HaloClassifier (purple) was trained exclusively on haloarchaeal genomes. Contigs within the 0.4–0.6 probability range are shown as “Unclassified”, distinguishing between true plasmid and true chromosomal fragments. Metrics shown include accuracy (overall correct predictions), Matthews correlation coefficient (MCC, balance between classes), precision (fraction of predicted positives that are correct), recall (fraction of true positives correctly predicted), and F1-score (harmonic mean of precision and recall).

Across all size ranges, HaloClassifier consistently outperformed the bacterial-focused tools in both accuracy and Matthews correlation coefficient (MCC). Notably, the high chromosome recall and plasmid precision observed for PlasFlow—and, to a lesser extent, PlasClass—can be misleading. These patterns are largely driven by a strong bias toward assigning fragments to the chromosome class: most contigs are classified as chromosomal, which inflates chromosome recall, while only a small fraction of true plasmid fragments are correctly identified as such, artificially increasing plasmid precision at the expense of plasmid recall.

For the Extralarge model, PlasFlow classified 762 fragments as chromosomal, of which 471 were truly chromosomal and 291 were plasmidic, resulting in high chromosome recall but low precision. Of the 500 plasmid fragments in the dataset, only 64 were predicted as plasmids—56 correctly and 8 incorrectly—which explains why PlasFlow exhibits such a high plasmid precision (0.8750) yet a very low plasmid recall (0.1614). The remaining 174 fragments fell below the 0.6 probability threshold for both classes and were labeled as unclassified; these comprised 21 chromosomal and 153 plasmid-derived fragments (17.4% of the dataset).

PlasClass showed a similar pattern but predicted more plasmids and left fewer fragments unclassified: 699 fragments were classified as chromosomal (419 correct, 280 plasmidic) and 261 as plasmidic (197 correct, 64 chromosomal), with only 4% of fragments (40) remaining unclassified. This led to a substantially higher plasmid recall than PlasFlow (0.4130 vs. 0.1614), while still retaining some misclassifications.

In sharp contrast, HaloClassifier showed a far more balanced and accurate behavior. For the same Extralarge dataset, it classified 487 fragments as chromosomal, correctly identifying 480 and misclassifying only 7 plasmid-derived fragments. Similarly, of the 485 fragments predicted as plasmids, 477 were correct and only 8 corresponded to chromosomal sequences. A total of 28 fragments (2.8%) remained unclassified—substantially fewer than in PlasFlow and comparable to PlasClass—split between 12 true chromosomal and 16 true plasmid fragments. These results in both chromosome and plasmid classes being recovered with high accuracy and minimal bias, reflecting the stability and reliability achieved by a model specifically trained on haloarchaeal genomes.

### Metagenomic performance assessment

To assess the performance of HaloClassifier on metagenomic assemblies, we analysed 233 haloarchaeal MAGs from the GEM dataset [33], comprising 57 enrichment-derived MAGs and 176 environmental MAGs. For each MAG, we calculated the proportion of its sequence classified by HaloClassifier at or above each of three confidence thresholds (0.6, 0.7 and 0.8). As shown in Figure 7A, increasing the confidence threshold resulted in fewer environmental MAGs having a high proportion of their sequence classified above that threshold. At a threshold of 0.6, two pronounced peaks appeared around 0.8 and 0.9, indicating that for most MAGs, 80–90% of their sequence was classified with ≥60% confidence. Increasing the threshold to 0.7 caused the second peak to shift downward to approximately 0.5, although a strong peak remained at 0.9. At the most stringent threshold (0.8), the distribution displayed three well-defined peaks: one near 0, a second around 0.4, and a slightly lower peak above 0.8. This pattern suggests that roughly one-third of the MAGs have more than 80% of their sequence classified with ≥80% confidence. For the enriched MAGs (Figure 7B), most genomes clustered at high values across all three thresholds. The complete prediction results for environmental and enrichment-derived MAGs are provided in Supplementary Files S20 and S21, respectively.

**Figure 7.**
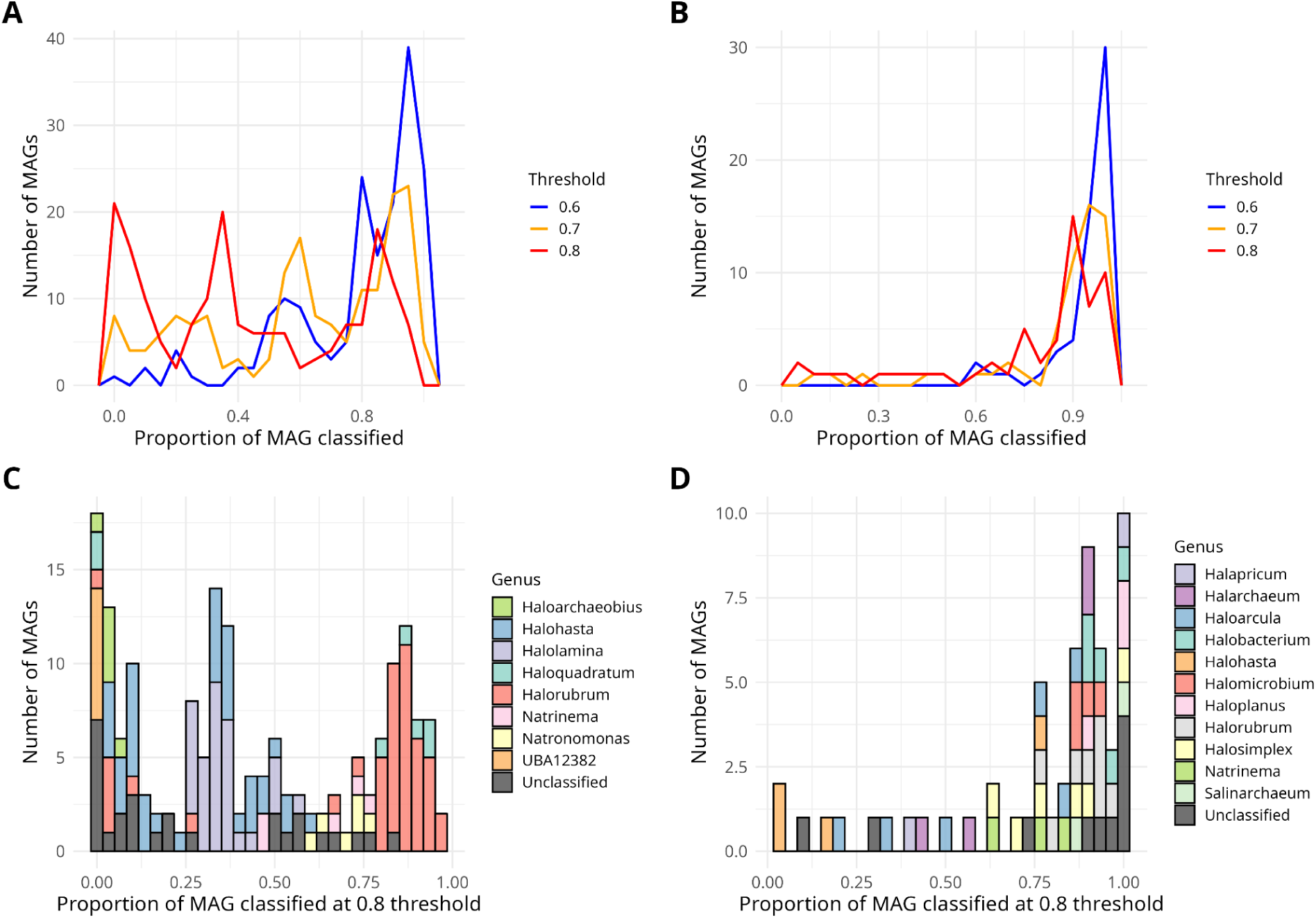
**Distribution of the proportion of each MAG’s sequence classified at or above three confidence thresholds**—0.6 (blue line), 0.7 (orange line), and 0.8 (red line)—for the environmental MAGs (**A**) and the enriched MAGs (**B**). Panels (**C**) and (**D**) show stacked histograms depicting the corresponding genus-level composition at the 0.8 threshold for environmental and enriched MAGs, respectively. In panel (**C**), the “Unclassified” category includes both MAGs that could not be assigned to any genus and those belonging to genera represented by fewer than five MAGs in the dataset; in panel (**D**), it includes only MAGs without a genus assignment. UBA12382 represents a *Hikarchaeia* lineage closely related to *Haloarchaea*, but its precise taxonomic relationship to *Haloarchaea* remains unresolved.

To further investigate the origin of the three-peak distribution observed at threshold 0.8 for non-enriched MAGs, we examined additional metadata features that might explain the pattern. Our initial hypothesis was that the sample origin could influence MAG quality and thereby classification performance. However, no meaningful relationship emerged (Figure S6). In contrast, when annotating the 0.8-threshold histogram with genus labels (Figure 7C), a clear association became evident. *Halorubrum* MAGs tended to exhibit higher proportions of confidently classified sequence, whereas other genera—such as *Haloarchaeobius*—showed the opposite trend. Comparison with the genera represented in the training dataset (Figure 2) revealed that *Halorubrum* had substantial representation (nine complete genomes), whereas *Haloarchaeobius* was entirely absent.

Interestingly, the genus *Haloquadratum*, also absent from the training set, showed strong classification performance: five of the seven *Haloquadratum* MAGs in the GEM database had more than 80% of their sequence confidently classified (≥80% confidence). *Halolamina* and *Halohasta*, likewise unrepresented in the training set, displayed intermediate values clustering around 0.4. The placeholder genus UBA12382, on the other hand, represents a *Hikarchaeia* lineage closely related to *Haloarchaea*, but whose precise taxonomic relationship to *Haloarchaea* remains unresolved [35,36]. Its low classification performance may therefore reflect its distinct evolutionary position relative to the haloarchaeal lineages represented in the training dataset.

In contrast, differences between genera were not apparent for the enriched MAGs. As shown in Figure 7D, most of these MAGs exhibited high proportions of confidently classified sequence at the 0.8 threshold, highlighting the influence of sample quality on downstream classification. Notably, the genus *Salinarchaeum*, absent from the training dataset, showed strong performance: the two *Salinarchaeum* MAGs in the GEM dataset were classified with ≥80% confidence across 87% and 100% of their sequence, respectively. Remarkably, the family to which this genus belongs—*Natronoarchaeaceae*—was also absent from the training dataset.

To further assess whether classification performance was associated with the phylogenetic relatedness between MAGs and the genomes represented in the training dataset, we used two complementary approaches. First, we assessed genomic relatedness using Average Nucleotide Identity (ANI), comparing each MAG with the genomes in the training dataset and retaining the highest ANI as the best-matching reference. We then assessed the relationship between this ANI and the proportion of MAG sequence classified with ≥80% confidence (Figure S7). Of the 176 environmental MAGs, 16 were excluded from the ANI analysis because their assemblies did not meet the quality requirements for reliable ANI estimation, leaving 160 MAGs. Environmental MAGs showed a strong positive association between ANI and classification performance (Spearman’s ρ = 0.796, *p* < 2.2 × 10⁻¹⁶; N = 160), with more similar MAGs tending to have a larger proportion of their sequence confidently classified. This relationship remained strong when ANI was weighted by the fraction of MAG sequence mapped to the reference genome (ρ = 0.772, *p* < 2.2 × 10⁻¹⁶; N = 160), indicating that the association was not driven solely by high ANI values supported by a small fraction of the MAG. In contrast, neither raw ANI (ρ = 0.057, *p* = 0.673; N = 57) nor mapping-weighted ANI (ρ = 0.096, *p* = 0.478; N = 57) was associated with classification performance in enriched MAGs.

As a complementary analysis, we then examined whether this pattern was also reflected in taxonomic relatedness (Table S4; Figure S8). Environmental MAGs showed markedly lower classification performance with increasing taxonomic distance: the median proportion of sequence classified with ≥80% confidence decreased from 0.727 for same-genus comparisons to 0.080 for same-family and 0.131 for different-family comparisons. Median ANI likewise decreased from 82.30% to 77.48% and 78.32%, respectively. However, the same-family category was relatively small (N = 6), and these results should therefore be interpreted cautiously. In contrast, enriched MAGs retained high classification performance across the three taxonomic categories, with median values of 0.881, 0.735, and 0.895 for same-genus, same-family, and different-family comparisons, respectively, despite decreasing median ANI values of 86.77%, 83.92%, and 79.86%. The same-family and different-family categories were both small (N = 4), further limiting the strength of these comparisons. Overall, despite the uneven sample sizes across taxonomic categories, these results support the ANI-based analysis, indicating a pronounced association between genomic relatedness and classification performance in environmental MAGs, but no clear association in enriched MAGs.

Additional analyses of the plasmid content of these MAGs, both in enriched and non-enriched datasets, were performed and are provided in the Supplementary Material (Figure S9; Table S5). Overall, plasmid-associated sequences represented a relatively small fraction of most MAGs, although substantial variation was observed between datasets. Enrichment-derived MAGs showed a median plasmid content of 0% (mean 3.9%), indicating that most genomes contained little or no plasmid-associated sequence. In contrast, environmental MAGs exhibited a significantly higher median plasmid content of 4.4% (mean 12.2%; Wilcoxon rank-sum test, *p* < 0.001). Consistent with this pattern, only 5.3% of enrichment-derived MAGs contained more than 20% plasmid-associated sequence, compared with 18.8% of environmental MAGs.

## Discussion

Genomic signatures (non-coding features), particularly *k*-mer composition and GC content, have dominated plasmid/chromosome classification approaches, owing to their strong predictive performance and relatively low computational cost [10,14,37,38]. However, the difficulty of reliably classifying short contigs [39], particularly those below 1 kb, has motivated the development of approaches that exploit sequence information in alternative ways, including graph-based methods that leverage *k*-mer connectivity [11] and, more recently, representation-learning approaches based on learned sequence embeddings [13]. To the best of our knowledge, coding-derived features—i.e., features explicitly derived from predicted coding regions or their properties—have received little attention as an additional source of information for distinguishing plasmid- and chromosome-derived sequences. This may partly reflect the substantially larger feature space offered by *k*-mer profiles, as well as the additional computational cost associated with gene prediction, which is required to derive coding-related features. Here, we show that, at least for *Haloarchaea*, coding-derived features can provide a viable and computationally tractable source of predictive information: despite comprising a much smaller feature space than *k*-mers (approximately 130 coding-derived features versus nearly 11,000 features for canonical *k*-mers of length 2–7), coding-derived features retained substantial predictive power and, in many cases, performed comparably to models based exclusively on non-coding features. Moreover, combining coding- and non-coding features significantly improved accuracy compared with either reduced-feature configuration, indicating that the two feature classes provide complementary sources of information.

The feasibility of incorporating coding-derived information is further supported by the feature-selection strategy implemented in HaloClassifier. Previous plasmid/chromosome classification approaches based on *k*-mer profiles have typically used the complete set of candidate *k*-mers as input features [10,14,37]. In contrast, our program incorporates a feature-selection step prior to model training, in which a parsimonious subset of informative predictors is identified. Combined with the use of four size-specific models, this approach allows the predictor set to adapt to the amount of sequence information available in each contig-length range while reducing computational complexity and runtime. Moreover, this strategy allowed us to examine the relative contribution of coding- and non-coding features across size ranges. Coding features accounted for 26%, 25%, 20%, and 11% of the predictors retained in the Small, Medium, Large, and Extralarge models, respectively, and for 30%, 26%, 21%, and 17% of the top 100 predictors ranked by Random Forest importance. Notably, in the Small model, the Only Coding configuration preserved plasmid recall at essentially the same level as the Final Model (75.61% vs. 75.58%; *p* = 0.677), whereas the No Coding configuration showed a significant decrease relative to the Final Model (74.53%; *p* < 0.001). These results suggest that coding-derived information is particularly valuable for shorter contigs, whereas *k*-mer features become increasingly informative as more sequence context is available.

Given the increasing availability of metagenomic data, we also assessed the ability of HaloClassifier to classify MAGs and to generalize beyond the taxonomic diversity represented in the training dataset. In environmental MAGs, lower genomic similarity to the training dataset was associated with markedly reduced classification performance. Median ANI decreased from 82.30% for same-genus comparisons to 77.48% and 78.32% for same-family and different different-family comparisons, respectively, while the corresponding median proportions of sequence classified with ≥80% confidence decreased from 0.727 for same-genus MAGs to 0.080 and 0.131 for same-family and different-family MAGs, respectively. In contrast, enriched MAGs showed a markedly different pattern: median ANI decreased from 86.77% for same-genus comparisons to 83.92% and 79.86% for same-family and different-family comparisons, respectively, yet the median proportion of confidently classified sequence remained high (0.881, 0.735, and 0.895, respectively). Thus, in enriched MAGs, the decrease in genomic similarity was not accompanied by a corresponding loss of classification performance, in contrast to the clear association observed for environmental MAGs. However, the same-family category comprised only six MAGs in the environmental dataset and four in the enriched dataset, while the different-family category comprised 14 and four MAGs, respectively; these comparisons should therefore be interpreted cautiously, particularly for the smaller groups. Together, these results indicate that taxonomic representation in the training dataset contributes to classification performance, but does not fully determine it, as other factors—potentially including genome reconstruction quality—can influence the successful classification of previously unseen lineages. The high-confidence predictions obtained for the two *Salinarchaeum* MAGs, belonging to the previously unseen family *Natronoarchaeaceae*, and for several environmental MAGs assigned to genera absent from the training dataset, such as *Haloquadratum*, further support the ability of HaloClassifier to generalize to haloarchaeal lineages that were not represented in the training data.

Beyond their predictive power, an additional advantage of coding-derived features is their interpretability. Whereas *k*-mer composition is often difficult to relate directly to biological mechanisms, coding-related patterns can reveal differences in codon usage or other sequence properties associated with protein-coding regions. Notably, in *Haloarchaea*, the GCC codon, which encodes alanine, emerged as the top-ranking feature in both the Extralarge and Large submodels. Haloarchaeal genomes are characterized by pronounced codon-usage biases, including a preference for GC-rich codons [40], and alanine-encoding codons have previously been identified among the preferred codons in this lineage [41]. Moreover, differences in codon composition between haloarchaeal host genomes and their extrachromosomal elements have been reported, including differences involving alanine-encoding codons [41]. In this context, the prominent contribution of GCC in our models suggests that differences in alanine codon usage may contribute to the distinction between plasmid- and chromosome-derived sequences and could reflect underlying differences in the functional or metabolic roles of these replicons, potentially including processes related to alanine metabolism. Although this hypothesis would require dedicated functional analyses to test, it illustrates how coding-derived features can provide biologically interpretable signals that would be difficult to identify using a purely compositional framework.

Given the strong performance of coding-only models, it is worth noting—although this lies beyond the scope of the present study—that coding-derived features may also prove valuable for plasmid/chromosome classification in (meta)transcriptomic datasets, where sequences are composed predominantly of coding regions and compositional features developed for genomic DNA may not be directly applicable. To our knowledge, no existing tool has specifically addressed plasmid/chromosome classification from transcriptomic sequences, potentially reflecting the limited applicability of conventional sequence-composition approaches to this type of data. In this context, the performance of the Only Coding model is particularly encouraging, as it retained most of the predictive signal and achieved performance within 1% accuracy of the Final Model. While this result is likely influenced by the high coding density characteristic of haloarchaeal genomes, and may therefore not generalize to organisms with substantially lower coding densities, it nevertheless suggests that coding-derived information alone can capture a substantial proportion of the signal required for plasmid/chromosome classification. Exploring whether coding-derived features can be exploited to distinguish plasmid- and chromosome-derived transcripts therefore represents a promising direction for future methodological development.

Finally, we highlight a particularly valuable application of HaloClassifier for *Haloarchaea*. Members of this lineage undergo extensive homologous recombination and display high levels of genomic promiscuity, with large DNA segments frequently exchanged among closely—and occasionally distantly—related strains [20,42,43]. In this context, HaloClassifier offers a practical framework for detecting genomic regions that may have experienced recombination or replicon-to-replicon movement. For contigs larger than 105 kb, the tool automatically partitions the sequence and generates an independent prediction for each fragment. When applied to well-characterized replicons—either chromosomes or (mega)plasmids—these windowed predictions can be examined to identify segments whose current replicon assignment differs from that inferred from their local sequence properties. Such an approach enables the detection of chromosomal regions that may have been excised and subsequently recombined into plasmids, as well as plasmid-derived segments that may have been historically integrated into the chromosome, yet still retain their original replicon-specific signals, thereby providing a means to reconstruct the evolutionary history of haloarchaeal genomes.

Moreover, to address cases in which a strain acquires large genomic fragments from more distantly related taxa, a promising future extension of HaloClassifier would be to incorporate an independent estimate of the taxonomic origin of each fragment—analogous to the strategy implemented in PlasFlow. Combining plasmid/chromosome classification with taxonomic inference would enable not only the identification of recombined regions, but also the tracing of their likely donor lineages, offering a powerful framework for dissecting the complex mosaicism characteristic of haloarchaeal genomes.

## Conclusions

In this study, we developed HaloClassifier, a machine-learning approach for plasmid/chromosome classification specifically adapted to the genomic characteristics of *Haloarchaea*. By combining coding-derived and non-coding sequence features and training separate models across four contig-size ranges, HaloClassifier achieved robust classification performance while allowing the contribution of different feature types to be evaluated across different sequence lengths. Coding-derived features retained substantial predictive power despite representing a much smaller feature space than *k*-mer profiles, and their contribution was particularly relevant for shorter contigs. The combination of both feature classes further improved classification performance, supporting their complementary nature. The feature-selection strategy additionally provided a parsimonious set of informative predictors, reducing the computational burden while enabling the identification of biologically interpretable signals, including codon-usage patterns.

Evaluation on metagenome-assembled genomes showed that HaloClassifier can generalize beyond the taxonomic composition of the training dataset, although performance varied depending on the genomic similarity and taxonomic representation of the evaluated MAGs. The strong predictions obtained for several lineages not represented in the training data indicate that taxonomic novelty does not preclude reliable classification, while the reduced performance observed for some environmental MAGs highlights the importance of genome quality and training-set diversity for metagenomic applications.

Overall, HaloClassifier provides a complementary framework for plasmid/chromosome classification in *Haloarchaea* that combines predictive performance, computational tractability, and feature interpretability. Beyond genomic classification, the strong performance of coding-derived features suggests potential applications to (meta)transcriptomic data, while the window-based predictions generated for long contigs could facilitate the identification of genomic regions with atypical replicon signatures and contribute to the study of recombination and replicon-to-replicon DNA transfer in this highly recombinogenic lineage. Future extensions incorporating taxonomic inference at the fragment level could further enable the identification and tracing of putative donor lineages involved in the mosaic evolution of haloarchaeal genomes.

## Availability and requirements

Project name: HaloClassifier

Project home page: https://github.com/Lorena0105/HaloClassifier

Operating system(s): Platform independent

Programming language: Python ≥ 3.10

Other requirements: Prodigal v2.6.3; pandas ≥ 2.3.3 and < 3.0; scikit-learn ≥ 1.7.2 and < 1.8; joblib ≥ 1.5.2 and < 2.0; tqdm ≥ 4.67.1

License: MIT

Any restrictions to use by non-academics: None

## List of Abbreviations

ENC: Effective number of codons
HGT: Horizontal gene transfer
MAG: Metagenome-assembled genome
MCC\: Matthews correlation coefficient
RSCU: Relative synonymous codon usage

## Declarations

### Ethics approval and consent to participate Not applicable

Consent for publication Not applicable.

### Availability of data and materials

HaloClassifier is freely available on GitHub under the MIT license: https://github.com/Lorena0105/HaloClassifier. The datasets generated and analysed during the current study, together with the supplementary files supporting the analyses, are available in Zenodo at [10.5281/zenodo.22233815]. The accession numbers of the 155 chromosomal sequences and 510 plasmid sequences used to construct the dataset are provided in Supplementary Files S1 (*File_S1_155_chromosomes.csv*) and S2 (*File_S2_510_plasmids_PLSDB.csv*), respectively. The accession numbers of the MAGs used for the environmental and laboratory-enriched datasets are provided in Supplementary Files S5 (*File_S5_MAGs_lab_enrichment.tsv*) and S6 (*File_S6_MAGs_environmental.tsv*), respectively. Additional supplementary files provide the CheckM-based genome filtering results, taxonomic corrections, feature-selection results, ranked feature lists, and HaloClassifier predictions for the MAG datasets. All supplementary files are available in the Zenodo repository cited above. The supplementary figures and tables accompanying this article are provided as Supplementary Information.

### Authors’ contributions

LLR conceived the study, designed and developed the machine learning tool, performed the analyses, interpreted the results, and wrote the manuscript. FGC contributed to the design of the data analyses and critically reviewed the manuscript. AFS critically reviewed the manuscript. All authors read and approved the final manuscript.

## Acknowledgements

The computations and analyses were performed on the high-performance computing cluster Garnatxa at the Institute for Integrative Systems Biology (I2SysBio, UV-CSIC).

