## Supplementary figures and tables for "HaloClassifier: integrating coding-signature features and *k*-mer composition for plasmid-chromosome discrimination in haloarchaeal genomes"

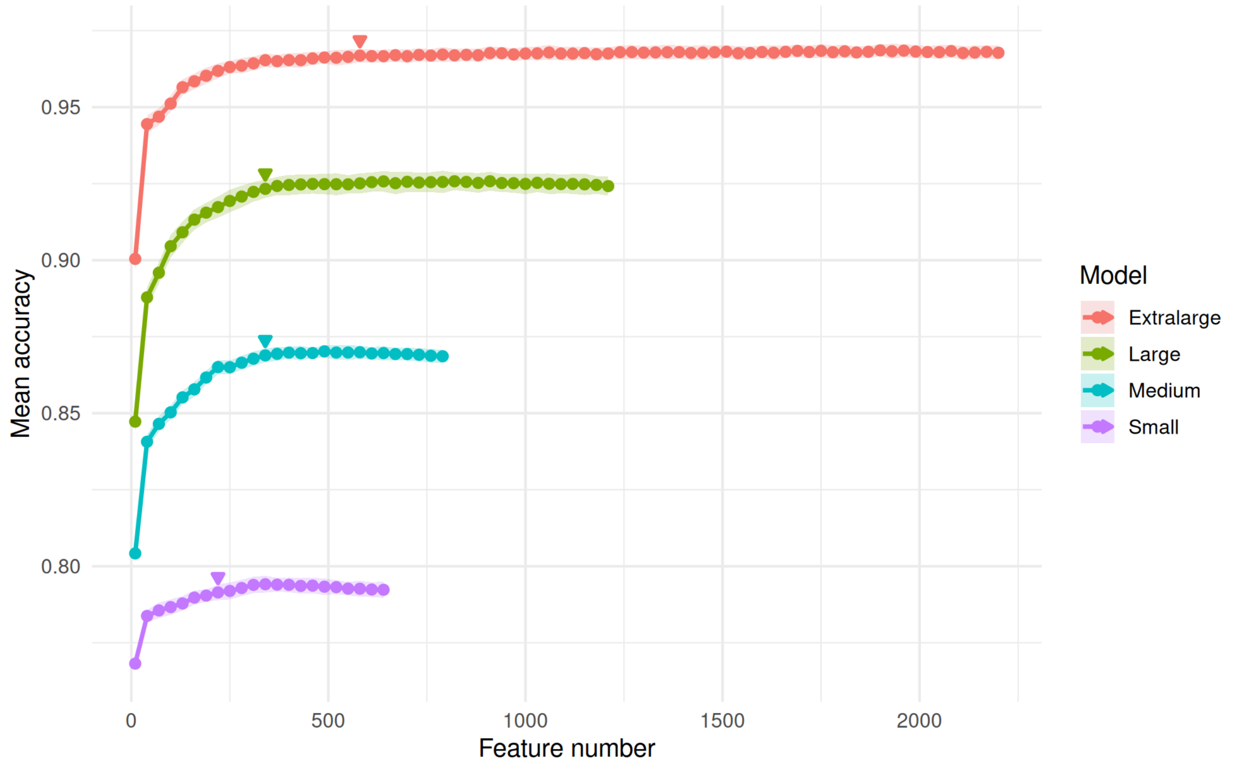


**Figure S1. Accuracy as a function of the number of variables tested during the coarse feature-selection phase for four different model configurations (Small, Medium, Large, Extralarge).** Shaded areas represent the standard deviation across repetitions. Colored arrows indicate the number of variables selected as optimal in the coarse phase, defined as the smallest set of variables whose mean accuracy falls within one standard deviation of the maximum mean accuracy. Early stopping was consistently triggered once the accuracy curve reached a stable maximum.

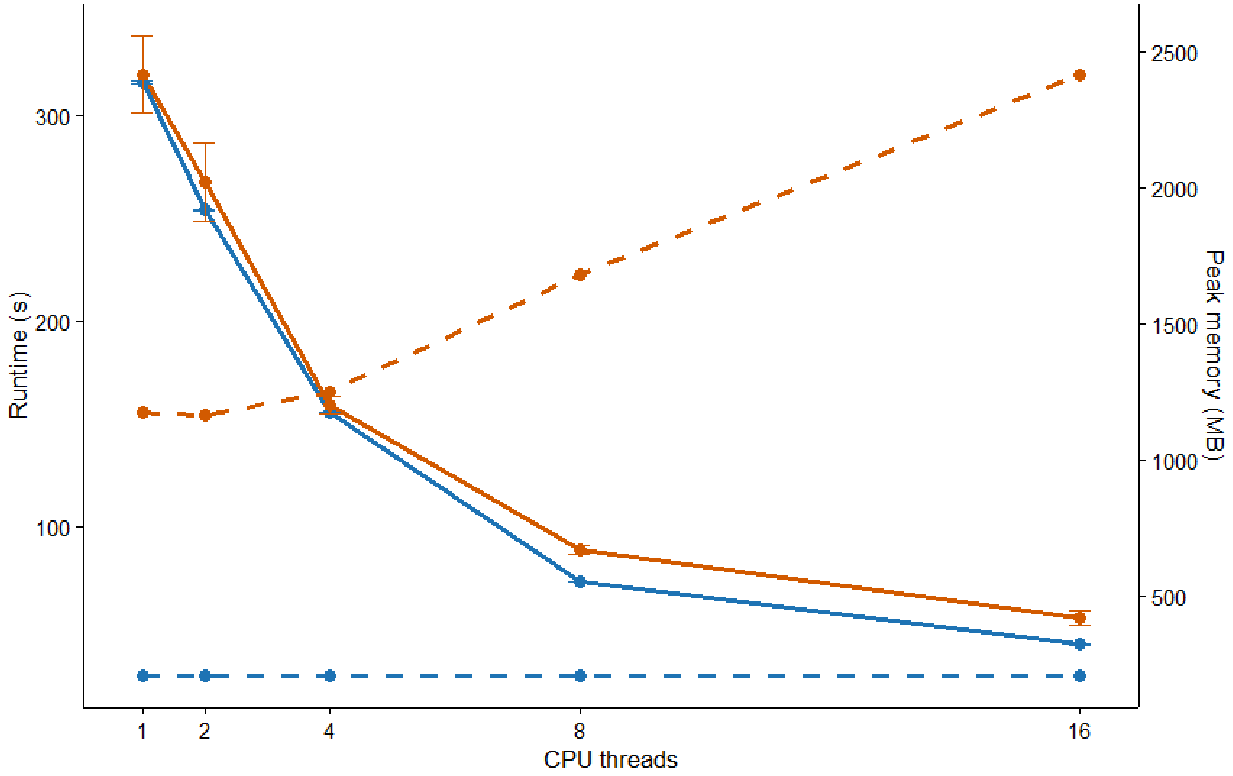


**Figure S2. Computational performance of HaloClassifier and PlasForest across different CPU thread counts.** Both pipelines were executed 10 independent times on a dataset of 1,000 randomly selected Extralarge contigs. Solid lines show runtime (seconds), whereas dashed lines show peak memory usage (MB). Error bars represent the standard deviation of the runtime measurements. HaloClassifier is shown in blue and PlasForest in orange.

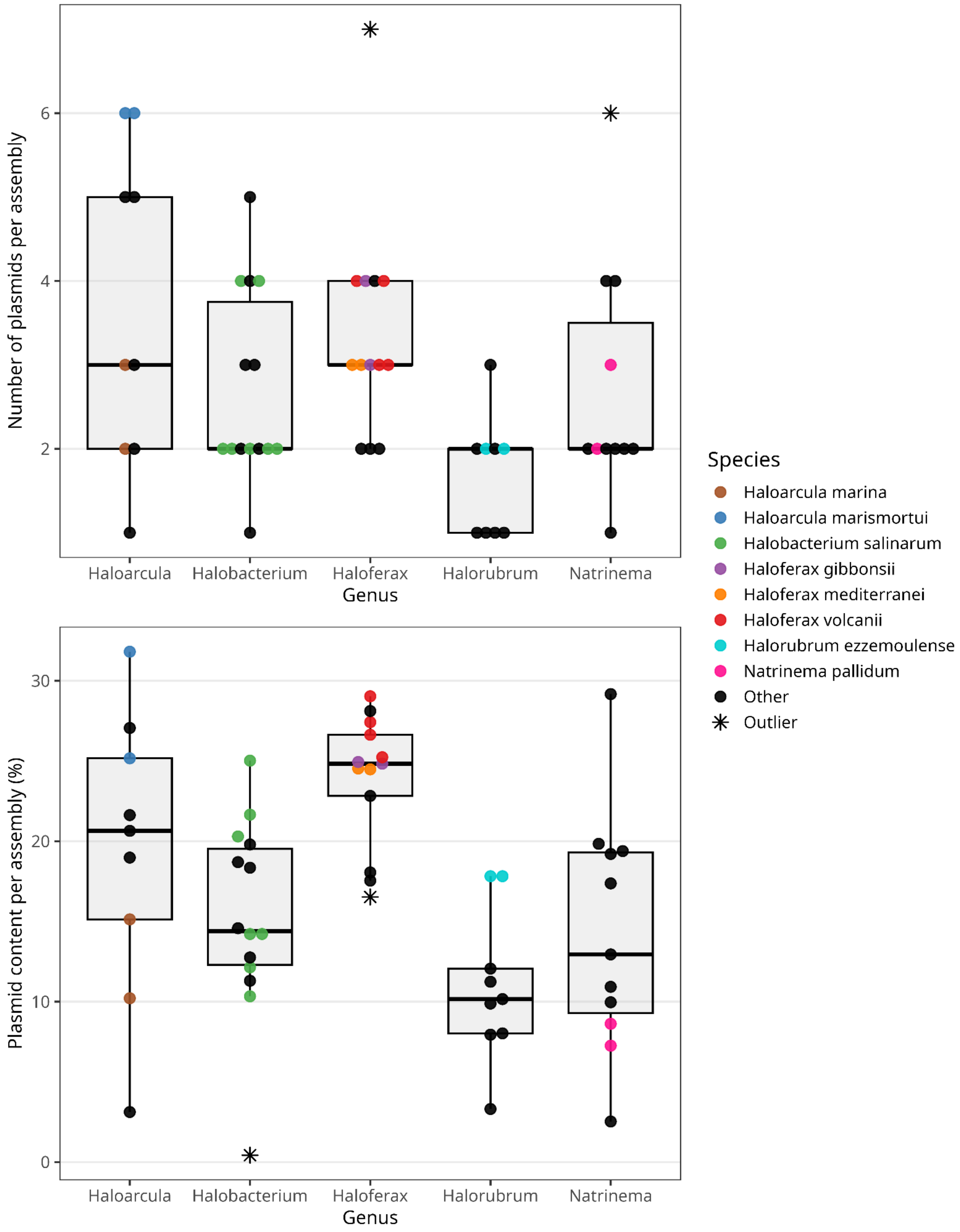


**Figure S3. Plasmid characteristics across selected *Haloarchaea* genera.** The top panel shows the distribution of the number of plasmids per assembly, represented as boxplots, whereas the bottom panel shows the distribution of plasmid content per assembly (% of total genomic DNA), also represented as boxplots. In both panels, species represented by two or more assemblies are colored individually, whereas singleton species or unclassified "sp." are shown in black. Outliers beyond 1.5 × IQR are highlighted as asterisks. The same color scheme and species labels apply to both panels.

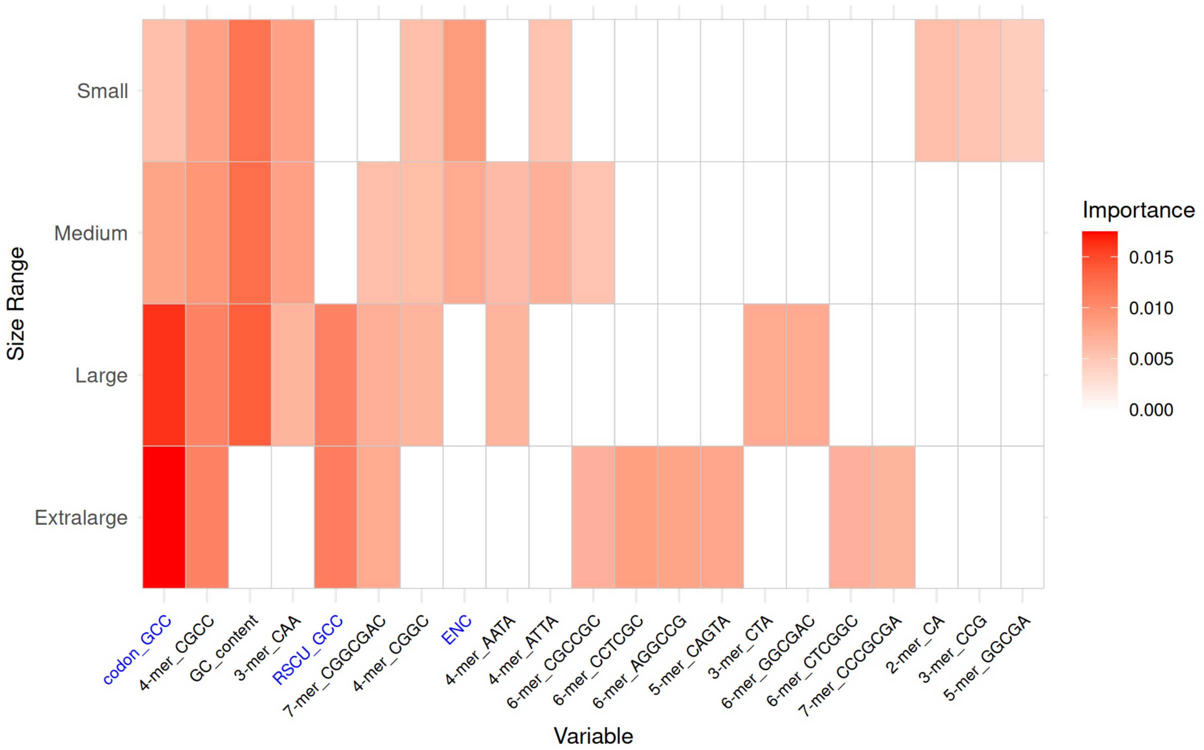


**Figure S4. Top 10 most important variables (Random Forest importance score) for each size-specific HaloClassifier model** (Extralarge, Large, Medium, and Small). Variables are arranged along the x-axis according to the highest importance score they achieve in any of the four models. Cell color intensity represents the relative importance of each variable within each model, with darker shades indicating higher importance. Coding-related features are highlighted in blue.

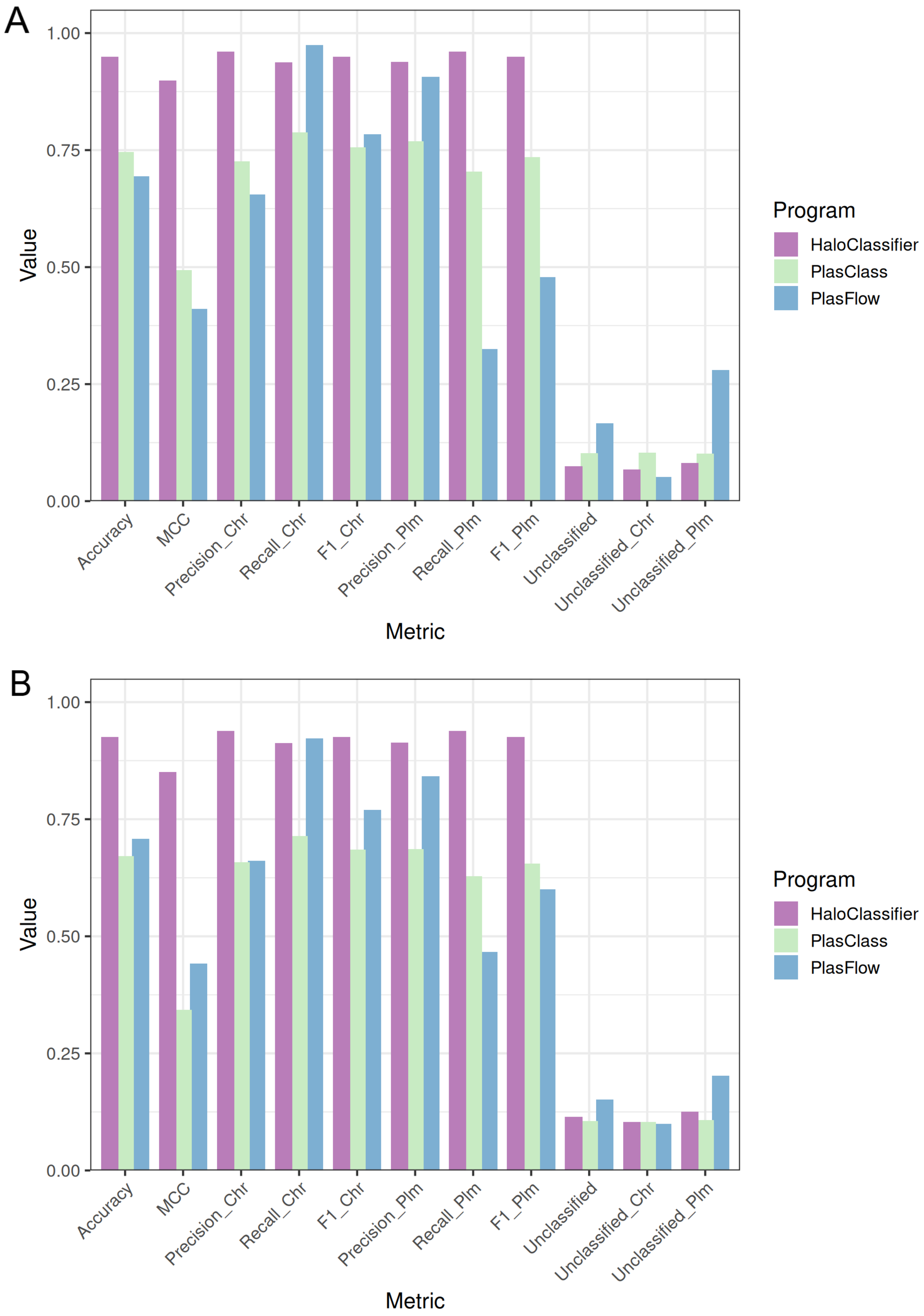
A
B
1.00
0.75
0.50
0.25
0.00
Value
Metric
Accuracy
MCC
Precision_Chr
Recall_Chr
F1_Chr
Precision_Plm
Recall_Plm
F1_Plm
Unclassified
Unclassified_Chr
Unclassified_Plm
Program
HaloClassifier
PlasClass
PlasFlow

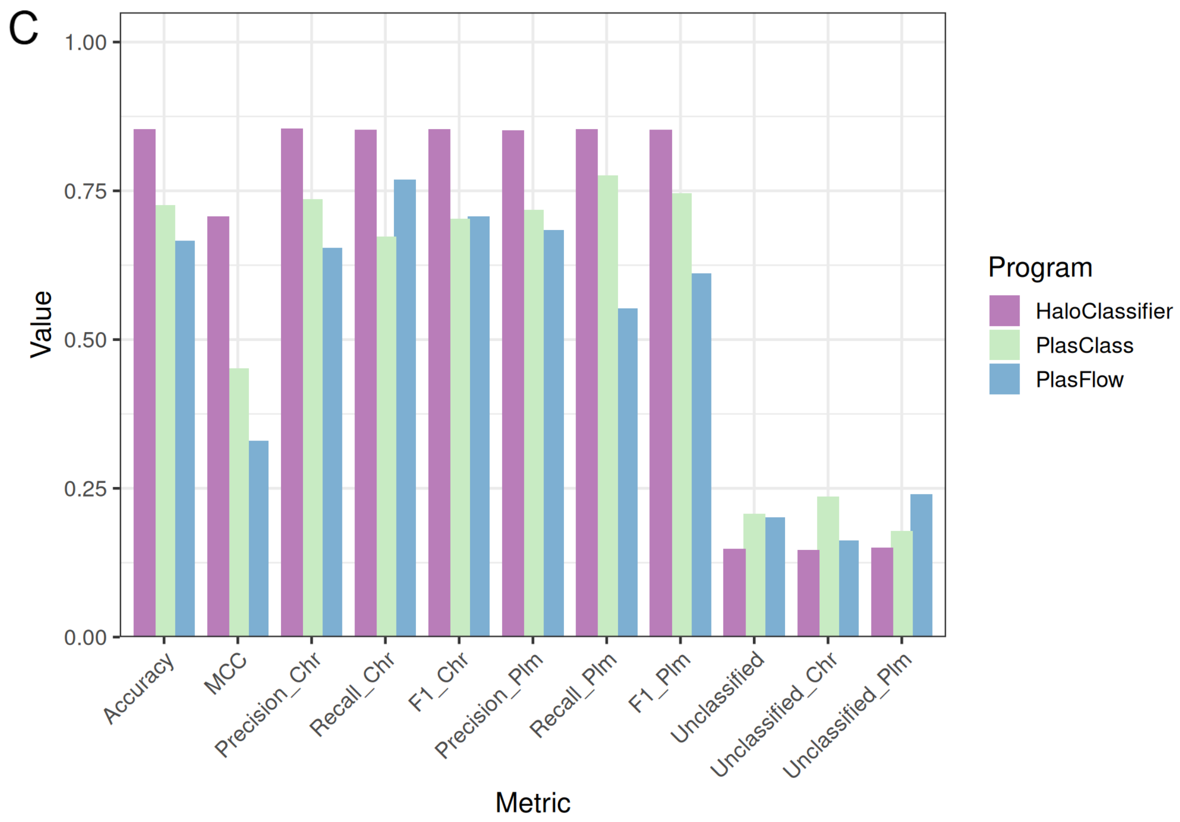


**Figure S5. Performance comparison of three plasmid identification tools on simulated contigs from complete haloarchaeal genomes across the Large (A), Medium (B), and Small (C) models.** PlasClass (green) and PlasFlow (blue) were developed using bacterial genomes, whereas HaloClassifier (purple) was developed specifically for haloarchaeal genomes. Contigs with predicted probabilities between 0.4 and 0.6 are shown as Unclassified, distinguishing between true plasmid and true chromosomal fragments. Performance metrics include accuracy (overall correct predictions), Matthews correlation coefficient (MCC; balanced performance across classes), precision (fraction of predicted plasmids that are correct), recall (fraction of true plasmids correctly identified), and F1-score (harmonic mean of precision and recall).

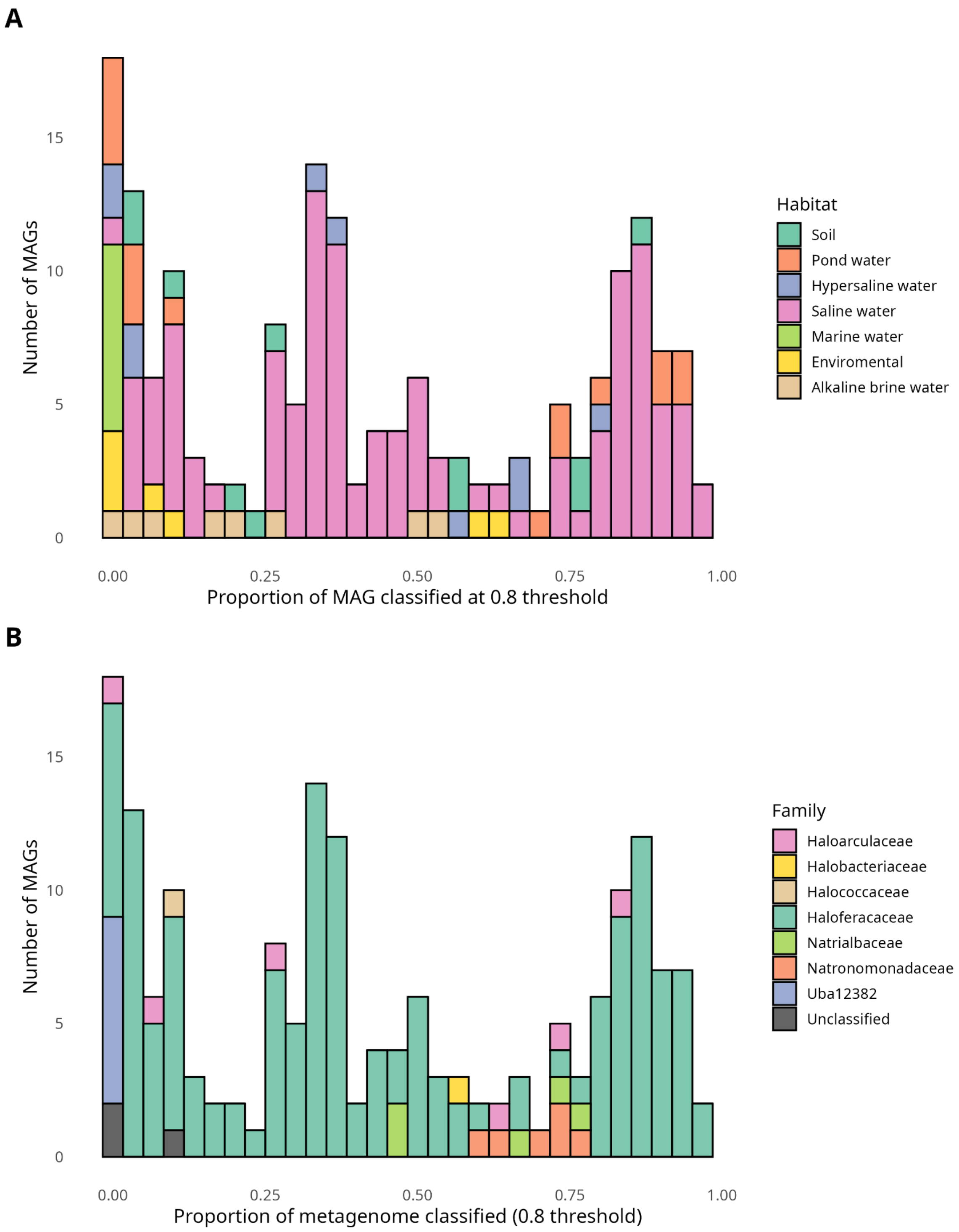


**Figure S6. Stacked histograms showing the distribution of environmental MAGs** according to the proportion of their assembled sequence classified with a probability ≥0.8. Panel A shows the corresponding environmental source, whereas panel B shows the family-level taxonomic composition. In B, the Unclassified category includes MAGs that could not be assigned to any haloarchaeal family.

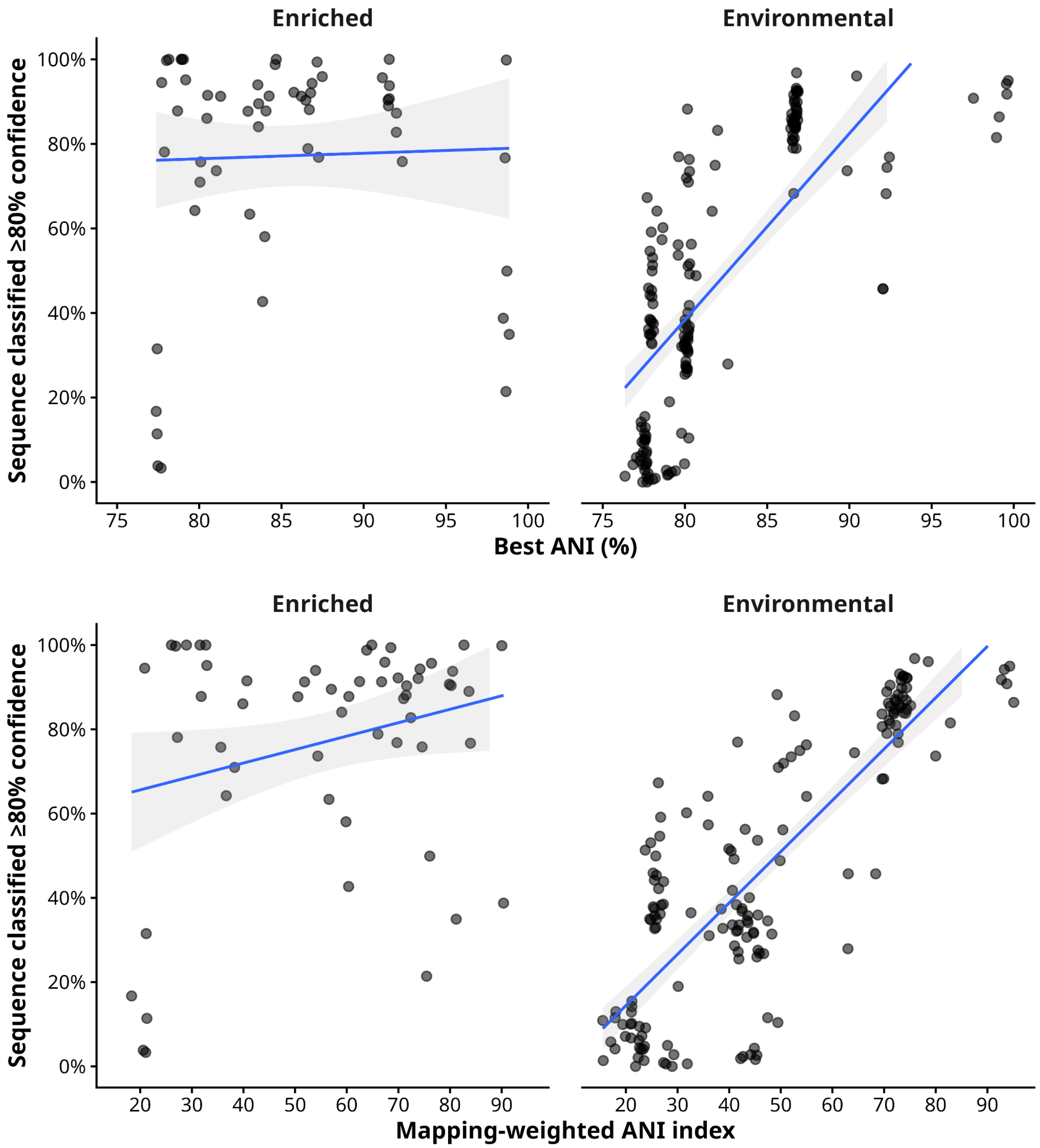


**Figure S7. Relationship between ANI and the proportion of MAG sequence classified with ≥80% confidence**. The top panel shows the relationship between the best ANI and the proportion of sequence classified with ≥80% confidence, whereas the bottom panel shows the corresponding relationship using a mapping-weighted ANI index, which accounts for both the ANI of the best-matching training genome and the fraction of MAG sequence mapped to the reference. Each point represents one MAG, and lines represent linear regressions with 95% confidence intervals. Spearman's rank correlation for the top panel was $\rho = 0.796$ ($p < 2.2 \times 10^{-16}$; N = 160) for environmental MAGs and $\rho = 0.057$ ($p = 0.673$; N = 57) for enriched MAGs. For the bottom panel, Spearman's $\rho$ was 0.772 ($p < 2.2 \times 10^{-16}$; N = 160) for environmental MAGs and 0.096 ($p = 0.478$; N = 57) for enriched MAGs.

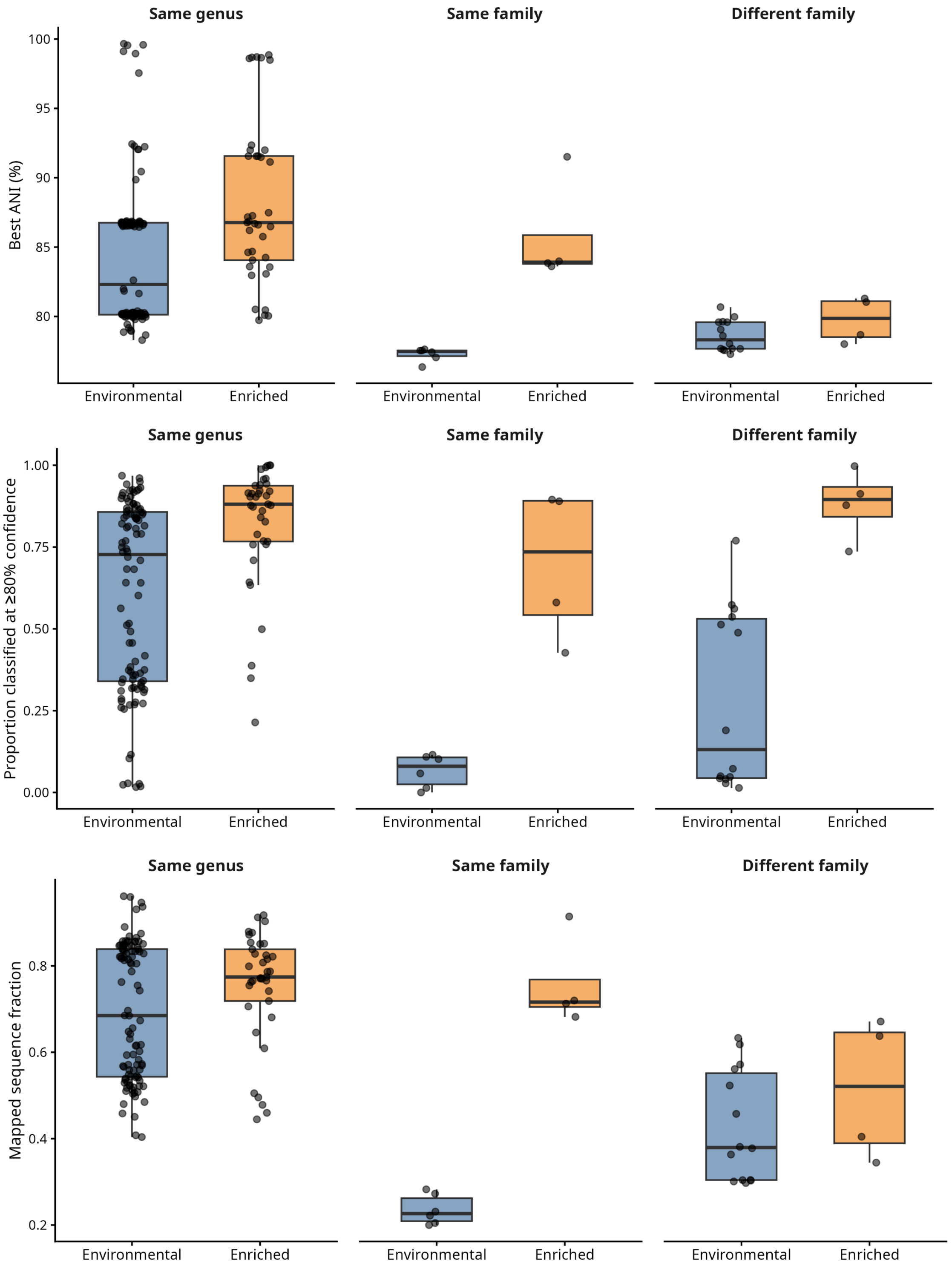


**Figure S8. Distribution of (top) best ANI, (middle) proportion of MAG sequence classified with ≥80% confidence, and (bottom) mapped fragment fraction** across environmental and enriched MAGs according to taxonomic relationship with the best-matching genome from the training dataset. Boxes indicate the interquartile range and median; black points represent individual MAGs.

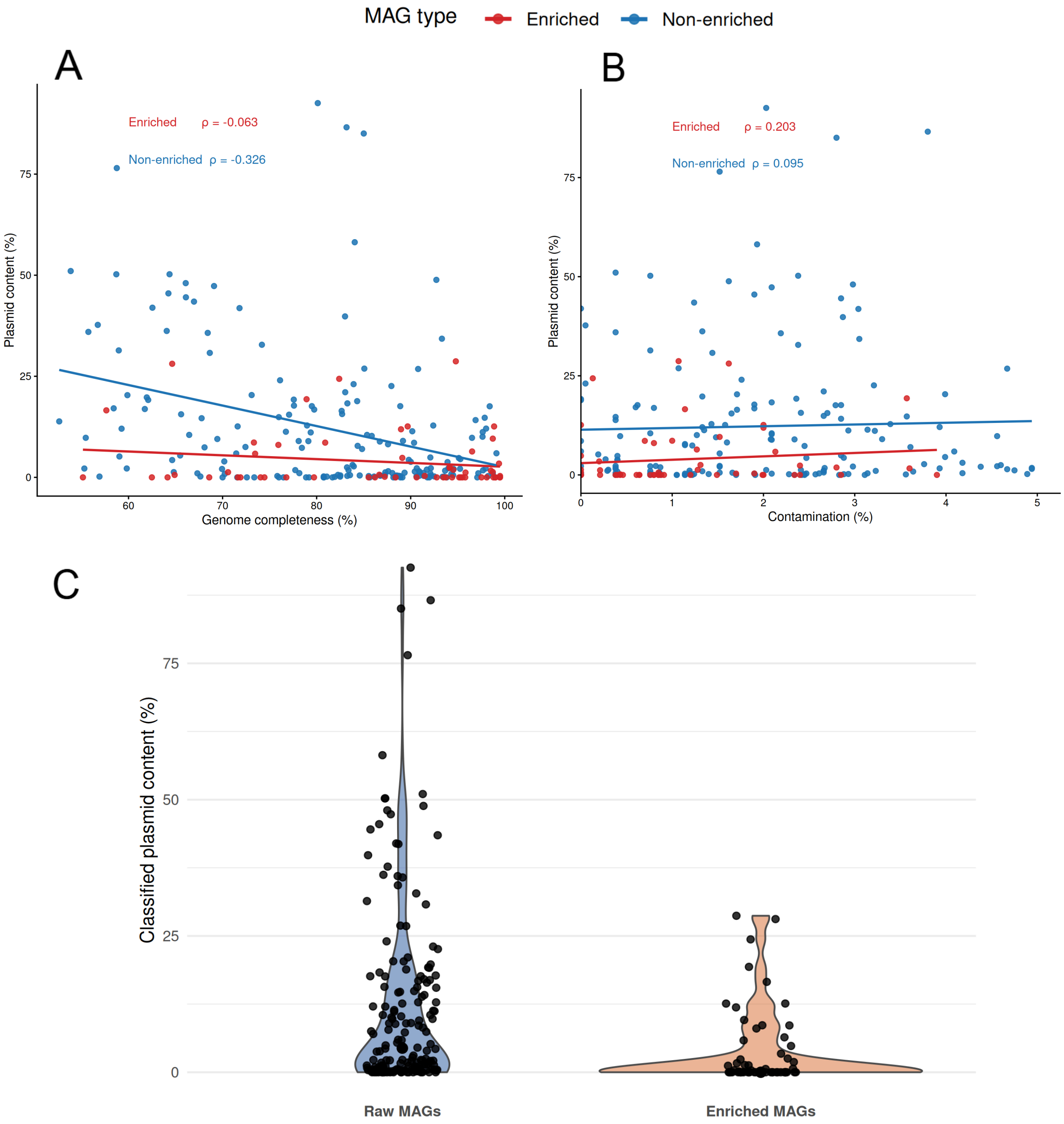


**Figure S9. Complementary analyses of plasmid content in haloarchaeal MAGs**. (**A**) Relationship between plasmid content (% of the assembled sequence classified as plasmid by HaloClassifier) and MAG completeness. (**B**) Relationship between plasmid content (%) and MAG contamination. In both panels, non-enriched and enriched MAGs are shown in blue and red, respectively. Solid lines represent linear regressions, and the corresponding slope coefficients are indicated. (**C**) Distribution of the proportion of plasmid-derived sequences within the moderate-confidence fraction of each MAG. For each genome, only sequences classified with a prediction probability ≥0.6 were considered, and the plotted value corresponds to the percentage of this fraction classified as plasmid.

**Table S1**. Performance comparison between the Final Model and the two reduced-feature configurations (Only Coding and No Coding) across the four contig-length categories. Panels correspond to (A) Extralarge, (B) Large, (C) Medium and (D) Small.

A. Extralarge

| Metric | Final Model (FM) | Only Coding (COD) | No Coding (NC) | COD vs FM | NC vs FM |
|---|---|---|---|---|---|
| Accuracy | 0.9668 ± 0.0024 | 0.9615 ± 0.0029 | 0.9652 ± 0.0025 | <0.001*** | <0.001*** |
| MCC | 0.9340 ± 0.0048 | 0.9233 ± 0.0057 | 0.9308 ± 0.0050 | <0.001*** | <0.001*** |
| Precision Chr | 0.9812 ± 0.0024 | 0.9739 ± 0.0025 | 0.9789 ± 0.0026 | <0.001*** | <0.001*** |
| Recall Chr | 0.9520 ± 0.0048 | 0.9486 ± 0.0051 | 0.9512 ± 0.0045 | <0.001*** | 0.106 |
| F1 Chr | 0.9663 ± 0.0025 | 0.9611 ± 0.0030 | 0.9648 ± 0.0026 | <0.001*** | <0.001*** |
| Precision Plm | 0.9532 ± 0.0045 | 0.9497 ± 0.0047 | 0.9523 ± 0.0042 | <0.001*** | 0.0462* |
| Recall Plm | 0.9816 ± 0.0024 | 0.9744 ± 0.0025 | 0.9794 ± 0.0026 | <0.001*** | <0.001*** |
| F1 Plm | 0.9672 ± 0.0023 | 0.9619 ± 0.0028 | 0.9656 ± 0.0025 | <0.001*** | <0.001*** |
| Num features | 553.8 ± 75.6263 | 122.6 ± 11.1915 | 594.8 ± 85.5677 | | |

B. Large

| Metric | Final Model (FM) | Only Coding (COD) | No Coding (NC) | COD vs FM | NC vs FM |
|---|---|---|---|---|---|
| Accuracy | 0.9230 ± 0.0026 | 0.9159 ± 0.0028 | 0.9169 ± 0.0024 | <0.001*** | <0.001*** |
| MCC | 0.8462 ± 0.0052 | 0.8321 ± 0.0055 | 0.8339 ± 0.0049 | <0.001*** | <0.001*** |
| Precision Chr | 0.9322 ± 0.0029 | 0.9257 ± 0.0031 | 0.9213 ± 0.0032 | <0.001*** | <0.001*** |
| Recall Chr | 0.9123 ± 0.0042 | 0.9045 ± 0.0047 | 0.9116 ± 0.0038 | <0.001*** | 0.162 |
| F1 Chr | 0.9221 ± 0.0027 | 0.9150 ± 0.0029 | 0.9165 ± 0.0025 | <0.001*** | <0.001*** |
| Precision Plm | 0.9142 ± 0.0038 | 0.9067 ± 0.0042 | 0.9126 ± 0.0034 | <0.001*** | 0.00579** |
| Recall Plm | 0.9337 ± 0.0030 | 0.9274 ± 0.0033 | 0.9222 ± 0.0035 | <0.001*** | <0.001*** |
| F1 Plm | 0.9238 ± 0.0026 | 0.9169 ± 0.0027 | 0.9174 ± 0.0024 | <0.001*** | <0.001*** |
| Num features | 420.6 ± 48.9966 | 119.8 ± 10.3562 | 422.2 ± 42.5450 | | |

C. Medium

| Metric | Final Model (FM) | Only Coding (COD) | No Coding (NC) | COD vs FM | NC vs FM |
|---|---|---|---|---|---|
| Accuracy | 0.8691 ± 0.0024 | 0.8588 ± 0.0024 | 0.8615 ± 0.0026 | <0.001*** | <0.001*** |
| MCC | 0.7383 ± 0.0048 | 0.7176 ± 0.0048 | 0.7232 ± 0.0052 | <0.001*** | <0.001*** |
| Precision Chr | 0.8639 ± 0.0028 | 0.8531 ± 0.0030 | 0.8522 ± 0.0030 | <0.001*** | <0.001*** |
| Recall Chr | 0.8759 ± 0.0028 | 0.8663 ± 0.0027 | 0.8743 ± 0.0031 | <0.001*** | <0.001*** |
| F1 Chr | 0.8699 ± 0.0024 | 0.8597 ± 0.0023 | 0.8631 ± 0.0026 | <0.001*** | <0.001*** |
| Precision Plm | 0.8745 ± 0.0027 | 0.8645 ± 0.0025 | 0.8712 ± 0.0030 | <0.001*** | <0.001*** |
| Recall Plm | 0.8623 ± 0.0031 | 0.8512 ± 0.0035 | 0.8487 ± 0.0035 | <0.001*** | <0.001*** |
| F1 Plm | 0.8683 ± 0.0025 | 0.8578 ± 0.0025 | 0.8598 ± 0.0027 | <0.001*** | <0.001*** |
| Num features | 358.6 ± 27.7458 | 106.0 ± 5.7735 | 323.2 ± 27.9090 | | |

D. Small

| Metric | Final Model (FM) | Only Coding (COD) | No Coding (NC) | COD vs FM | NC vs FM |
|---|---|---|---|---|---|
| Accuracy | 0.7941 ± 0.0029 | 0.7897 ± 0.0026 | 0.7883 ± 0.0027 | <0.001*** | <0.001*** |
| MCC | 0.5901 ± 0.0059 | 0.5808 ± 0.0052 | 0.5789 ± 0.0054 | <0.001*** | <0.001*** |
| Precision Chr | 0.7726 ± 0.0027 | 0.7709 ± 0.0026 | 0.7649 ± 0.0027 | 0.0013** | <0.001*** |
| Recall Chr | 0.8326 ± 0.0042 | 0.8234 ± 0.0037 | 0.8315 ± 0.0038 | <0.001*** | 0.0757 |
| F1 Chr | 0.8015 ± 0.0029 | 0.7963 ± 0.0025 | 0.7968 ± 0.0026 | <0.001*** | <0.001*** |
| Precision Plm | 0.8192 ± 0.0039 | 0.8112 ± 0.0034 | 0.8161 ± 0.0035 | <0.001*** | <0.001*** |
| Recall Plm | 0.7558 ± 0.0036 | 0.7561 ± 0.0035 | 0.7453 ± 0.0037 | 0.677 | <0.001*** |
| F1 Plm | 0.7862 ± 0.0031 | 0.7827 ± 0.0027 | 0.7791 ± 0.0029 | <0.001*** | <0.001*** |
| Num features | 259.4 ± 33.6749 | 99.2 ± 13.3604 | 219.0 ± 26.6930 | | |

**Table S2.** Confusion matrices for HaloClassifier, PlasFlow and PlasClass at the probability threshold used for benchmarking (0.6). Panels correspond to (A) Extralarge, (B) Large, (C) Medium and (D) Small contig-length categories. For each tool, the predicted class is shown in the rows and the true class in the columns. Contigs with predicted probabilities between 0.4 and 0.6 are reported as Unclassified.

A. Extralarge

| Tool | Predicted class | Chromosome | Plasmid | Total |
|---|---|---|---|---|
| HaloClassifier | Chromosome | 480 | 7 | 487 |
| | Unclassified | 12 | 16 | 28 |
| | Plasmid | 8 | 477 | 485 |
| | Total | 500 | 500 | 1000 |
| PlasFlow | Chromosome | 471 | 291 | 762 |
| | Unclassified | 21 | 153 | 174 |
| | Plasmid | 8 | 56 | 64 |
| | Total | 500 | 500 | 1000 |
| PlasClass | Chromosome | 419 | 280 | 699 |
| | Unclassified | 17 | 23 | 40 |
| | Plasmid | 64 | 197 | 261 |
| | Total | 500 | 500 | 1000 |

B. Large

| Tool | Predicted class | Chromosome | Plasmid | Total |
|---|---|---|---|---|
| HaloClassifier | Chromosome | 437 | 18 | 455 |
| | Unclassified | 34 | 41 | 75 |
| | Plasmid | 29 | 441 | 470 |
| | Total | 500 | 500 | 1000 |
| PlasFlow | Chromosome | 462 | 243 | 705 |
| | Unclassified | 26 | 140 | 166 |
| | Plasmid | 12 | 117 | 129 |
| | Total | 500 | 500 | 1000 |
| PlasClass | Chromosome | 353 | 133 | 486 |
| | Unclassified | 52 | 51 | 103 |
| | Plasmid | 95 | 316 | 411 |
| | Total | 500 | 500 | 1000 |

C. Medium

| Tool | Predicted class | Chromosome | Plasmid | Total |
|---|---|---|---|---|
| HaloClassifier | Chromosome | 409 | 27 | 436 |
| | Unclassified | 52 | 63 | 115 |
| | Plasmid | 39 | 410 | 449 |
| | Total | 500 | 500 | 1000 |
| PlasFlow | Chromosome | 415 | 213 | 628 |
| | Unclassified | 50 | 101 | 151 |
| | Plasmid | 35 | 186 | 221 |
| | Total | 500 | 500 | 1000 |
| PlasClass | Chromosome | 320 | 166 | 486 |
| | Unclassified | 52 | 54 | 106 |
| | Plasmid | 128 | 280 | 408 |
| | Total | 500 | 500 | 1000 |

D. Small

| Tool | Predicted class | Chromosome | Plasmid | Total |
|---|---|---|---|---|
| HaloClassifier | Chromosome | 364 | 62 | 426 |
| | Unclassified | 73 | 75 | 148 |
| | Plasmid | 63 | 363 | 426 |
| | Total | 500 | 500 | 1000 |
| PlasFlow | Chromosome | 322 | 170 | 492 |
| | Unclassified | 81 | 120 | 201 |
| | Plasmid | 97 | 210 | 307 |
| | Total | 500 | 500 | 1000 |
| PlasClass | Chromosome | 257 | 92 | 349 |
| | Unclassified | 118 | 89 | 207 |
| | Plasmid | 125 | 319 | 444 |
| | Total | 500 | 500 | 1000 |

**Table S3**. Performance of HaloClassifier across different probability thresholds (0.6–0.9) evaluated on the independent 20% test set for the four contig-length categories. Accuracy was calculated considering only classified fragments, whereas the percentage of unclassified fragments and the number of classified fragments relative to the total test set are also reported.

| Contig size | Threshold | Accuracy (classified) | Unclassified (%) | Classified / Total |
|---|---|---|---|---|
| Extralarge | 0.6 | 0.9806 | 3.55 | 5631 / 5838 |
| Extralarge | 0.7 | 0.9878 | 8.86 | 5321 / 5838 |
| Extralarge | 0.8 | 0.9927 | 18.23 | 4774 / 5838 |
| Extralarge | 0.9 | 0.9962 | 37.08 | 3673 / 5838 |
| Large | 0.6 | 0.953 | 8.69 | 9736 / 10662 |
| Large | 0.7 | 0.9711 | 18.96 | 8640 / 10662 |
| Large | 0.8 | 0.9812 | 31.99 | 7251 / 10662 |
| Large | 0.9 | 0.9884 | 51.37 | 5185 / 10662 |
| Medium | 0.6 | 0.9121 | 12.55 | 18999 / 21725 |
| Medium | 0.7 | 0.9453 | 26.59 | 15948 / 21725 |
| Medium | 0.8 | 0.9659 | 43.23 | 12333 / 21725 |
| Medium | 0.9 | 0.9826 | 67.15 | 7136 / 21725 |
| Small | 0.6 | 0.8442 | 17.07 | 18528 / 22343 |
| Small | 0.7 | 0.8868 | 36.56 | 14174 / 22343 |
| Small | 0.8 | 0.9201 | 57.27 | 9547 / 22343 |
| Small | 0.9 | 0.9526 | 79.97 | 4476 / 22343 |

**Table S4. Classification performance and ANI according to the taxonomic relationship between MAGs and their best-matching training genomes.** Median classification refers to the proportion of MAG sequences classified with ≥80% confidence. IQR corresponds to the interquartile range of this proportion. Median ANI represents the percentage nucleotide identity between the MAG and its best-matching genome from the training dataset. N indicates the number of MAGs in each group. *Unknown* indicates cases in which the taxonomic relationship could not be unambiguously determined because required family or genus information was unavailable. Of the 176 environmental MAGs, 16 were excluded from the ANI analysis because their assemblies did not meet the quality requirements for reliable ANI estimation; all 16 belonged to the *unknown* category.

| Dataset | Relationship | N | Median_classification | IQR_classification | Median_ANI |
|---|---|---|---|---|---|
| Environmental | Same genus | 100 | 0.727 | 0.517 | 82.30 |
| Environmental | Same family | 6 | 0.080 | 0.082 | 77.48 |
| Environmental | Different family | 14 | 0.131 | 0.487 | 78.32 |
| Environmental | Unknown | 40 | 0.130 | 0.347 | 77.81 |
| Enriched | Same genus | 37 | 0.881 | 0.171 | 86.77 |
| Enriched | Same family | 4 | 0.735 | 0.349 | 83.92 |
| Enriched | Different family | 4 | 0.895 | 0.091 | 79.86 |
| Enriched | Unknown | 12 | 0.863 | 0.846 | 77.80 |

**Table S5**. Summary statistics of plasmid-associated sequence content detected by HaloClassifier in metagenome-assembled genomes (MAGs) from the GEM dataset. Enrichment-derived and environmental MAGs are reported separately. Plasmid content is expressed as the percentage of the total assembled genome classified as plasmid (probability threshold ≥ 0.6). Mean, standard deviation (SD), median, quartiles (Q1 and Q3), minimum and maximum values are shown, together with the percentage of MAGs containing more than 10% or more than 20% plasmid-associated sequence. The plasmid content distributions differed significantly between enrichment-derived and environmental MAGs (Wilcoxon rank-sum test, $p = 5.88 \times 10^{-7}$).

| Dataset | Mean | SD | Median | Q1 | Q3 | Min | Max | >10% | >20% |
|---|---|---|---|---|---|---|---|---|---|
| Enrichment | 3.93 | 7.16 | 0 | 0 | 4.82 | 0 | 28.7 | 14 | 5.3 |
| Environmental | 12.24 | 17.68 | 4.42 | 0.47 | 16.81 | 0 | 92.56 | 38.1 | 18.8 |
